# A point mutation in *Ttc26* causes lumbar spinal cord fusion and synchronous hind-limb locomotion in *hop* mice

**DOI:** 10.1101/2020.05.13.093799

**Authors:** Nadine Bernhardt, Fatima Memic, Anna Velica, Michelle A. Tran, Jennifer Vieillard, Taha Chersa, Leif Andersson, Patrick J. Whelan, Henrik Boije, Klas Kullander

## Abstract

Identifying the spinal circuits controlling locomotion is critical for unravelling the mechanisms controlling the production of gaits. Development of the circuits governing left-right coordination relies on axon guidance molecules such as ephrins and netrins. To date, no other class of proteins have been shown to play a role during this process. Here we have analyzed *hop* mice, which walk with a characteristic hopping gait using their hind legs in synchrony. Fictive locomotion experiments suggest that a local defect in the ventral spinal cord contributes to the aberrant locomotor phenotype. *Hop* mutant spinal cords had severe morphological defects, including the absence of the ventral midline and a poorly defined border between white and grey matter. The *hop* mice represent the first model where the left and right central pattern generators (CPGs) are fused to form one central CPG, with a synchronous gait as a functional consequence. These defects were exclusively found in the lumbar domain and were associated with abnormal developmental processes, including a misplaced notochord and reduced induction of ventral progenitor domains. While the underlying mutation in *hop* mice has been suggested to lie within *Ttc26*, other genes in close vicinity have been associated with gait defects. By replicating the point mutation within *Ttc26*, employing CRISPR technology, we observed mice with an identical phenotype, thereby verifying the hop mutation. Thus, we show that the assembly of the lumbar CPG network is dependent on a fully functional TTC26 protein.

## Introduction

In the early 20th century, Thomas Graham-Brown demonstrated that the basic pattern for stepping could be generated by the spinal cord without descending or peripheral input (1). Within his model of spinal locomotor control he implemented Sherrington’s term “half-center” for two groups of reciprocally organized neurons mutually inhibiting each other to provide a coordinated pattern for stepping. Such local spinal networks, capable of generating rhythmic motor output, are referred to as central pattern generators (CPGs). Studies on the spinal cord locomotor CPG have identified key principles for the development and function of neuronal circuitry. For example, mouse mutants with abnormal locomotor coordination have been informative for understanding CPG organization and function (2–16).

The CPGs controlling hindlimb muscle activity during locomotion are located in the ventral part of the lumbar spinal cord (17,18) whereas CPGs controlling forelimb activity are located in the cervical spinal cord. Each side of the midline contains CPGs capable of rhythm generation where commissural inhibitory and excitatory projections ensure left-right coordination. Inhibitory actions between CPGs on different rostrocaudal levels of the spinal cord coordinate ipsilateral flexor and extensor muscles. The coordination between the left and right side of the body is maintained by commissural interneurons (CINs) that project to the contralateral side of the cord and innervate interneurons and motor neurons (4,14,19–23). Left-right activity persists when the dorsal spinal cord is removed, whereas left-right alternating fictive locomotion degrades after cutting the spinal cord ventral commissure. These experiments illustrate that ventromedial commissural interneurons are critical for bilateral coordination (12,13,18).

Neuronal connections form during embryonic development when differentiating neurons send their axons, navigating through the embryonic environment to synaptic targets. This process is mediated by conserved families of axon guidance proteins including netrins, ephrins, slits and semaphorins (24) and impaired axon guidance in the spinal cord results in locomotor coordination dysfunction (2,3,10,25–27).

In the present study, we analyzed *hop* mice that walk with a characteristic hopping gait using the hind legs in synchrony. Mice carrying the hop-sterile (*hop* or hydrocephalic-polydactyly, *hpy)* mutation appeared spontaneously in 1967 in the C57BL/10J strain and the mutation was first found to be localized at the proximal end of chromosome 6 (28–31). This was later identified as a point mutation within the *Ttc26* gene, suggested to be responsible for impaired hedgehog signaling (32).

However, since other genes in the close vicinity have been associated with gait defects, it remains unclear whether the point mutation is causative for the *hop* locomotor phenotype. Using CRISPR technology we introduced a single point mutation within the *Ttc26* gene, which reproduced the anatomical and behavioral phenotype observed in *hop* mice. Further, we show that the synchronous gait in *hop* mice is the result of a CPG half-centre fusion within the lumbar domain of the spinal cord, caused by an abnormal notochordal Shh signaling during development, resulting in migration defects and an absent ventral midline.

## Results

### Aberrant synchronous hindlimb coordination in *hop* mice correlates to local spinal cord neuronal circuitry

Homozygous *hop* mutants show distinct phenotypes such as preaxial polydactyly of all feet (Fig. S1), sperm tail deficiency resulting in male sterility, and hydrocephalus. Although there have been no detailed reports on the locomotor pattern produced by *hop* mutant animals, it has been described that they exhibit a characteristic hopping gait (28,33). We confirmed and extended these observations by conducting gait analysis and electrophysiological experiments. Gait analysis revealed that during locomotion, mice homozygous for the *hop* mutation, herein referred to *hop* mice, alternated their forelimbs while their hindlimbs moved in synchrony (Fig. 1A, B). To assess the onset of this hopping pattern, we evoked locomotor activity by pinching the tail, followed by electromyographic (EMG) recordings from the *tibialis anterior* muscles in conscious early postnatal (P0-P5) *hop* and control animals (n=2, respectively). We found that *hop* mice produced an air locomotor pattern of synchronous hindlimb movements while the flexor/extensor activity in the hindlimbs was normal. As expected, control neonatal mice produced a typical left-right and flexor-extensor alternating pattern (Fig. 1C-E).

**Figure 1:**
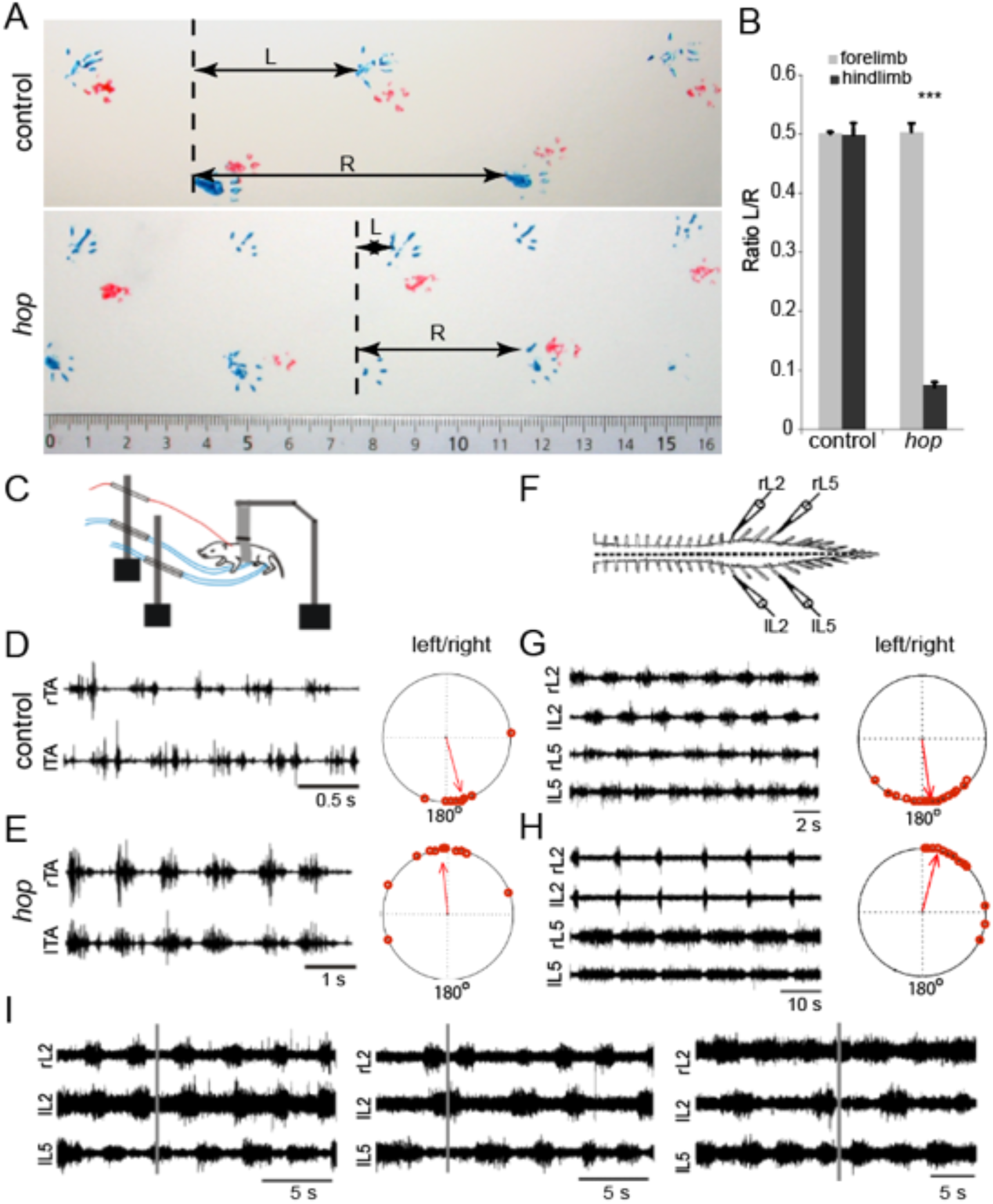
Abnormal gait in hind limbs of *hop* mice. (A) Gait analysis painting forelimbs (red) and hind limbs (blue) show that control mice move their fore- and hind limbs in an alternating pattern while *hop* mice move their hind limbs in a synchronous manner. Horizontal arrow L is the distance between left and right paw and R the distance covered by the same right paw. (B) Graph showing quantification of walking pattern with the ratio L/R (n=4 animals per group over a distance of 30 cm, p *** p < 0.001). (C-E) *hop* mice show a characteristic synchronous left-right air-stepping pattern *in vivo*. (C) Schematic drawing of experimental setup for recording from neonatal mice. (D-E) Raw EMG recordings from the left and right tibialis muscles of *hop* (E) and control (D) mice following a tail pinch stimulus illustrating representative synchronous and alternating traces, respectively. The circular plots in (D-E) represent the phase of onset of the left and right tibialis anterior bursts with respect to each other. Each dot represents an individual burst while the arrow indicates the tendency for the bursts to be coupled to each other. A phase value close to zero indicates synchrony in the case of the *hop* while a value close to 180° suggests an alternating pattern in the case of the control animals (P<0.01). (F-H) The *hop* deficit can be reproduced *in vitro* using isolated spinal cord preparations. (F) Schematic illustration of ventral root recording set-up measuring fictive locomotion. (G-H) Control mice exhibit an alternating (G) and *hop* mice a synchronous (H) left-right pattern. Circular plots in (G and H) indicating phase values across experiments. Each point equals one 60 second section of data (5 per animal). The length of the arrow provides an index of the strength of the coupling between the rhythms and the direction equals phase. (I) Bath application of GABA and glycinergic reuptake blockers affect the stability of the synchronous rhythm in the hop mutant mouse. Control rhythm in *hop* mice (top) evoked by bath application of NMDA, 5-HT and DA. 20 minutes after the addition of sarcosine and nipecotic acid (middle and right, respectively) one L2 root would double burst occasionally and at random.

To determine whether deficiencies in the spinal cord CPG could cause the locomotor phenotype, we performed experiments using isolated spinal cord preparations. Following bath application of 5-HT, DA and NMDA, we found that in the majority of *hop* mutants (20/28), a synchronous fictive locomotor rhythm developed between left-right ventral roots. In contrast, the alternating ipsilateral flexion-extension like pattern between the L2 and L5 roots was preserved (Fig. 1F-H). The data represent an average for 5 min following the development of a sustained rhythm (L2-L2: 1.10 ± 0.06 (PTCC), 5.61 ± 0.68 s (cycle period), 0.05 ± 0.01 (phase lag); L2-L5: -1.04 ± 0.06 (PTCC), 0.52 ± 0.03 (phase lag); n = 20). The synchronous fictive locomotion we observed in *hop* mice is in accordance with the *in vivo* gait analyses as well as the EMG recordings. However, the data indicate that the fictive locomotion in *hop* mice is less stable than in control mice. In animals that generated synchronous segmental patterns and ipsilateral alternation, we found that the ipsilateral pattern of alternation was maintained following hemisection, suggesting that the circuitry coordinating flexor-extensor alternation is intact in *hop* mutants (Fig. S2). To further establish whether synchronous left-right rhythmic activity was the dominant pattern, we used electrical stimulation of the *cauda equina* or dorsal roots to elicit a rhythm (34,35). We found that in *hop* mice, electrical stimulation did evoke a synchronous rhythm similar to that observed using pharmacological stimulation (Fig. S2C).

Previous studies have shown that neurotransmitter balance over the midline is critical for the left-right coordination output of the CPG, and during fictive locomotion, the balance between excitation and inhibition can be shifted using pharmacological strengthening (3,36–38). Therefore, we tested whether simultaneous bath application of glycine (sarcosine) and GABA (nipecotic acid or NO-711) reuptake blockers would alter the rhythm to an alternating pattern in *hop* mice. However, upon application of glycine/GABA reuptake blockers, the synchronous pattern did not switch to an alternating pattern but instead became less regular (Fig. 1I).

Taken together, our locomotor behavior analysis in *hop* mice confirmed the synchronous pattern previously reported and our *in vitro* analyses indicate that defects in the lumbar local spinal cord circuitry are sufficient to explain the altered hindlimb coordination. Further, changing the balance of excitatory and inhibitory connections over the midline did not shift the synchronous locomotor output of *hop* mice.

### Altered spinal cord morphology in *hop* mice

Since our electrophysiological experiments suggested that the aberrant locomotor phenotype arose from local spinal neuronal circuitry output, we next examined the morphology of the spinal cord. Gross morphology of wild type postnatal spinal cords is characterized by a cervical and lumbar enlargement, which contain, among others, the neurons that coordinate fore- and hindlimb movement, respectively. In *hop* mice, the size and length of the cervical enlargement was comparable to controls whereas the lumbar enlargement was hardly detectable (Fig. 2A-D). Further, wild type spinal cords had a clearly visible ventral spinal artery fed by a radicular artery termed the artery of Adamkiewicz, which in wild type mice usually originates at the level of L_1-2_ (Fig. 2C, arrow). In the lumbar region of *hop* spinal cords, the blood supply was evidently abnormal including the absence of a ventral spinal artery together with an undefined artery of Adamkiewicz (Fig. 2D).

**Figure 2:**
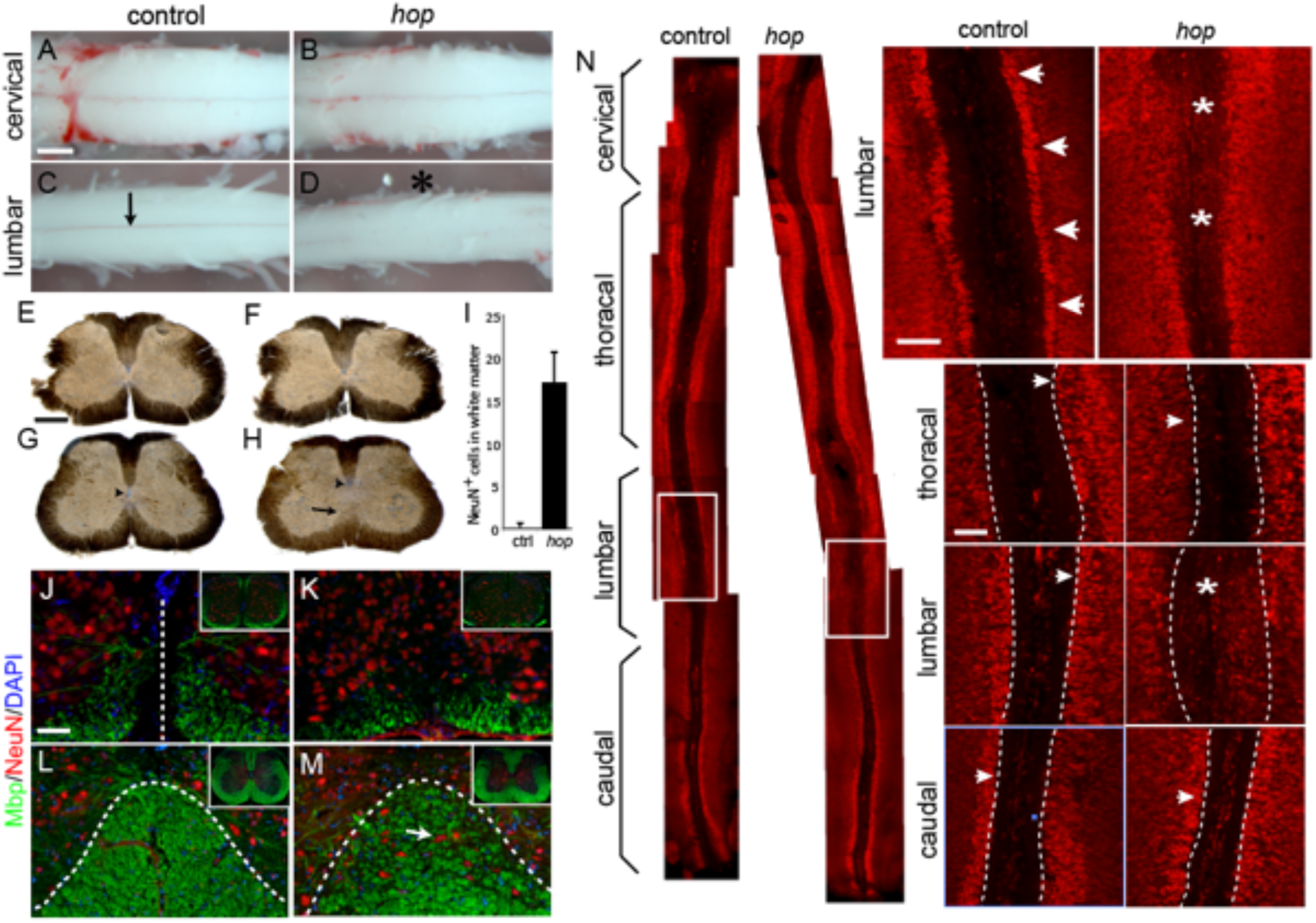
Altered spinal cord morphology in postnatal *hop* mice. (A-D) Photomicrographs of adult cervical and lumbar spinal cords. At cervical levels from control (A) and *hop* (B) mice, no difference is apparent. At lumbar levels, the lumbar enlargement of the spinal cord is less prominent in *hop* mice (D) compared to control mice (C). The clearly visible ventral spinal artery in control wild type mice (arrow in C) is absent from the lumbar part of the *hop* spinal cord (position noted as star in D). (E-H) Peroxidase staining on 60 μm transverse sections of control (E,G) and *hop* (F,H) spinal cords showing abnormal morphology in the lumbar spinal cord of *hop* mice. (H) The central canal is dislocated dorsally and is smaller in size (arrowhead), the ventral commissure is missing and the ventral white matter is disorganized or absent (black arrow). (I) Quantification of number NeuN positive cells/section (n=10 per genotype) in the lumbar white matter in the *hop* adult mice compared to control. (J-M) Immunohistochemistry staining with NeuN in red and MBP in green shows misplaced neurons in the grey matter of ventral spinal cord in *hop* mice at P5 (K) and adult (M) compared to control P5 (J) and adult (L). Dotted line in (J) indicates the midline, and in (L-M) the border between white and grey matter. (N) Photomicrographs of immunohistochemistry staining with NeuN in red on spinal cord open book preparations demonstrates lumbar specific defects in *hop* mice. White boxes indicate areas of higher magnification to the right. Arrows points to the border between white matter and NeuN positive cells and stars denote areas where aberrant NeuN positive cells are found. Scale bars 300 μm (A-D), 150 μm (E-H), and 50 μm (J-N).

Transverse sections of the cervical spinal cord from *hop* mice showed no morphological differences compared to controls (Fig. 2E, F). In contrast, transverse sections of lumbar spinal cords from *hop* mice revealed an altered morphology in the ventral spinal cord, including an undefined border between white and grey matter, the absence of a ventral funiculus and a dorsally shifted, poorly delineated central canal (Fig. 2H). Immunohistochemistry, using NeuN as a marker for neurons (grey matter) and myelin basic protein (MBP) as a marker for myelinated axons (white matter), showed less myelinated structures and misplaced neurons within the white matter in *hop* mutants (Fig. 2I-M).

Next, we used the spinal cord open book preparation, in which the spinal cord is cut dorsally along the rostrocaudal axis, and the left and right halves of the spinal cord are dissected to reveal the inside of the two halves and the remaining ventral part. In this preparation, we found that the lumbar part in *hop* mice was severely affected and had a diffuse border between the NeuN positive neurons and the ventral area, whereas the thoracic and caudal parts were unaffected (Fig. 2N).

Other mouse mutants with locomotor phenotypes have been reported to display severe defects in several brain regions including the corpus callosum, hippocampal-, anterior-, habenular- and posterior commissure as well as the pontine nucleus and cerebellar structures (2,39–43). However, morphological analysis of brains from *hop* mice, which have earlier been reported to display enlarged ventricles likely connected to the hydrocephalic phenotype, did not show any further brain abnormalities (Fig. S3).

Together, these findings suggest that the synchronous gait in *hop* mice is the result of ventral spinal cord alterations limited to the lumbar region controlling the hindlimbs. The rostro-caudal specificity is further supported by the observed forelimb alternating locomotion and normal cervical spinal cord morphology.

### Aberrant commissural fibres and neurotransmitter phenotype in *hop* mice

Cross-inhibitory and excitatory actions of commissural interneurons (CINs) between the two sides of the locomotor CPG ensure left-right coordination (18,44). In an effort to further explain the aberrant locomotor phenotype, we examined the neurotransmitter profile of the neurons interspersed in the ventral white matter of *hop* mice using *in situ* hybridization. GABA/glycinergic, glutamatergic and cholinergic neurons were identified by detecting mRNA encoding the vesicular inhibitory amino acid transporter (VIAAT), the vesicular glutamate transport (Vglut2), or the vesicular acetylcholine transporter (VAChT). We found both VIAAT and Vglut2 positive neurons in the ventral funiculus of *hop* mice demonstrating that these neurons can be either excitatory or inhibitory (Fig. 3A, B, D, E). Additionally, we found that central canal cholinergic cells were ventrally and medially displaced in *hop* mice compared to controls (Fig. 3C, F) whereas no apparent difference was found regarding position and size of motor neurons.

**Figure 3:**
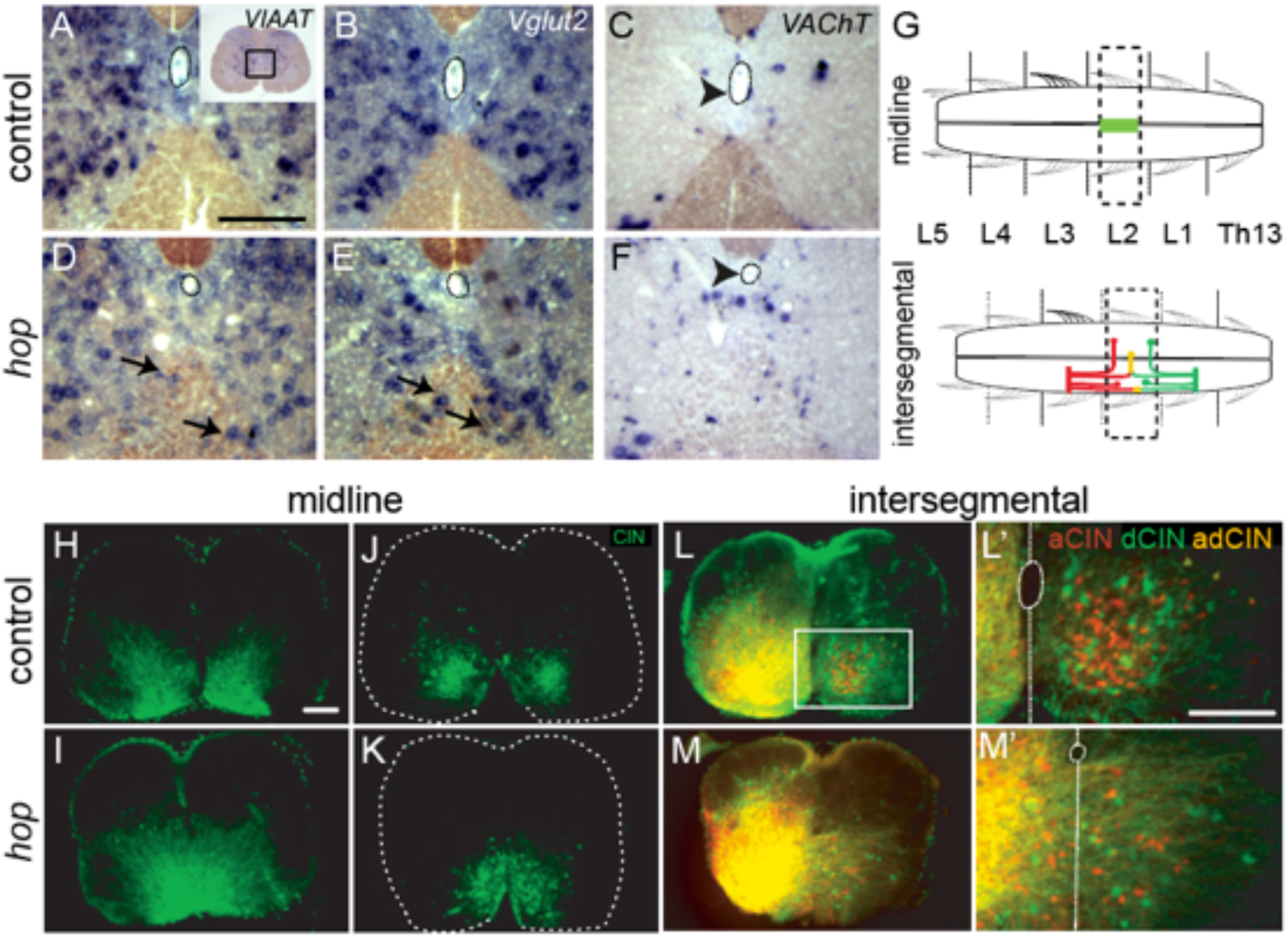
Neurotransmitter phenotype and aberrant commissural fibres in the ventral lumbar spinal cord of postnatal *hop* mice. (A-F) In situ hybridization on adult 60 μm floating sections using probes against VIAAT (A,D), Vglut2 (B,E) and VAChT (C,F). In control mice, neurons marked with either VIAAT (A) or Vglut2 (B) respect the border between white and grey matter whereas in *hop* mice, VIAAT (D) or Vglut2 (E) mRNA expressing neurons, intermingle with the white matter. VAChT stained cholinergic cells are found around the central canal in control (C) but are ventrally displaced relative to the central canal in *hop* mice (F). Black box in (A) indicates area of enlargements. Neurons in the white matter are marked with black arrows. Arrowheads indicate central canal. Scale bars 50μm (G-L). (G) Schematics of midline tracing using FDA to show local projecting commissural interneurons (H-K) or intersegmental FDA and RDA tracing to show ascending (aCINs), descending (dCINs) and bifurcating commissural interneurons (adCINs) in (L-M). (H, I) Transverse section within the L1 level of spinal cords traced on L2 shows that the fibres freely cross the midline in *hop* (I) compared to control mice (H). (J, K) Transverse section within the L2 level of spinal cords traced on L2 shows a lower number of commissural neurons, which are dispersed over the midline in *hop* mice (K) but not in controls (J). (L-M) Aberrant fibres cross over the midline in spinal cords of *hop* mice (M-M’) ventral from the central canal and the amount of aCINs, dCINs and adCINs are reduced compared to controls (L-L’). White box in (L) indicates the area of close-up (L’, M’). The outline of the spinal cord is indicated by a dashed line (J, K), central canal and midline are indicated by a dotted line (L’, M’). Scale bars (H-M) 100μm and (L’-M’) 200μm.

Ventromedial CINs with axons crossing in the ventral commissure are necessary and sufficient for left-right coordination (18), which prompted us to investigate the CINs in *hop* mice. Short- and long-range projecting CINs were examined by tracer applications of fluorescein dextran amine (FDA) and rhodamine dextran amine (RDA) in P0 mice (Fig. 3G). Midline retrograde tracing using FDA to identify locally short projecting CINs, revealed that CINs at the lumbar level 1 (L1, Fig. 3H, I) and the cell bodies positioned at lumbar level 2 (L2, Fig. 3J, K) cluster together on the ventral midline of *hop* mice compared to the two separate clusters of CINs seen in control mice. To study the long-range projecting CINs, we examined three different subpopulations; the intersegmental ascending (RDA), descending (FDA), and bifurcating (RDA/FDA) CINs in both lumbar and cervical spinal cord. In control mice, our tracings experiments showed that the three different populations were all in the ventromedial area, whereas in *hop* mice, the CINs were diffusely dispersed and partly located on the midline (Fig. 3L, M; Fig. S4).

Taken together, these results show that the ventromedial inhibitory and excitatory CINs responsible for normal left-right coordination are clustered together on the ventral midline in the *hop* mice resulting in a loss of ventral midline separating the two CPGs on each side.

### An early developmental defect underlies the altered functionality in *hop* mice

CPG activity in rodents develops between embryonic day (E)12 and birth. Correct assembly and maturation results in an established left-right coordination circuitry just before birth (45–47). The prominent ventral spinal cord phenotype observed in postnatal *hop* mice prompted us to investigate the onset of the morphological phenotype. Immunohistochemistry on E15.5 *hop* spinal cord tissue using microtubule associated protein 2 (Map2), to visualize fibers, and NeuN, to visualize neurons, revealed a similar fused spinal cord phenotype as seen in the adult (Fig. 4A-D). In addition, using RDA tracing at E15.5 and E12.5, we found that the traceable CIN population in the *hop* embryos were misplaced and had aberrant fibers crossing the midline (Fig. 4E-H). These data indicate deviant ventral spinal cord formation likely due to interrupted developmental processes including midline axon guidance.

**Figure 4:**
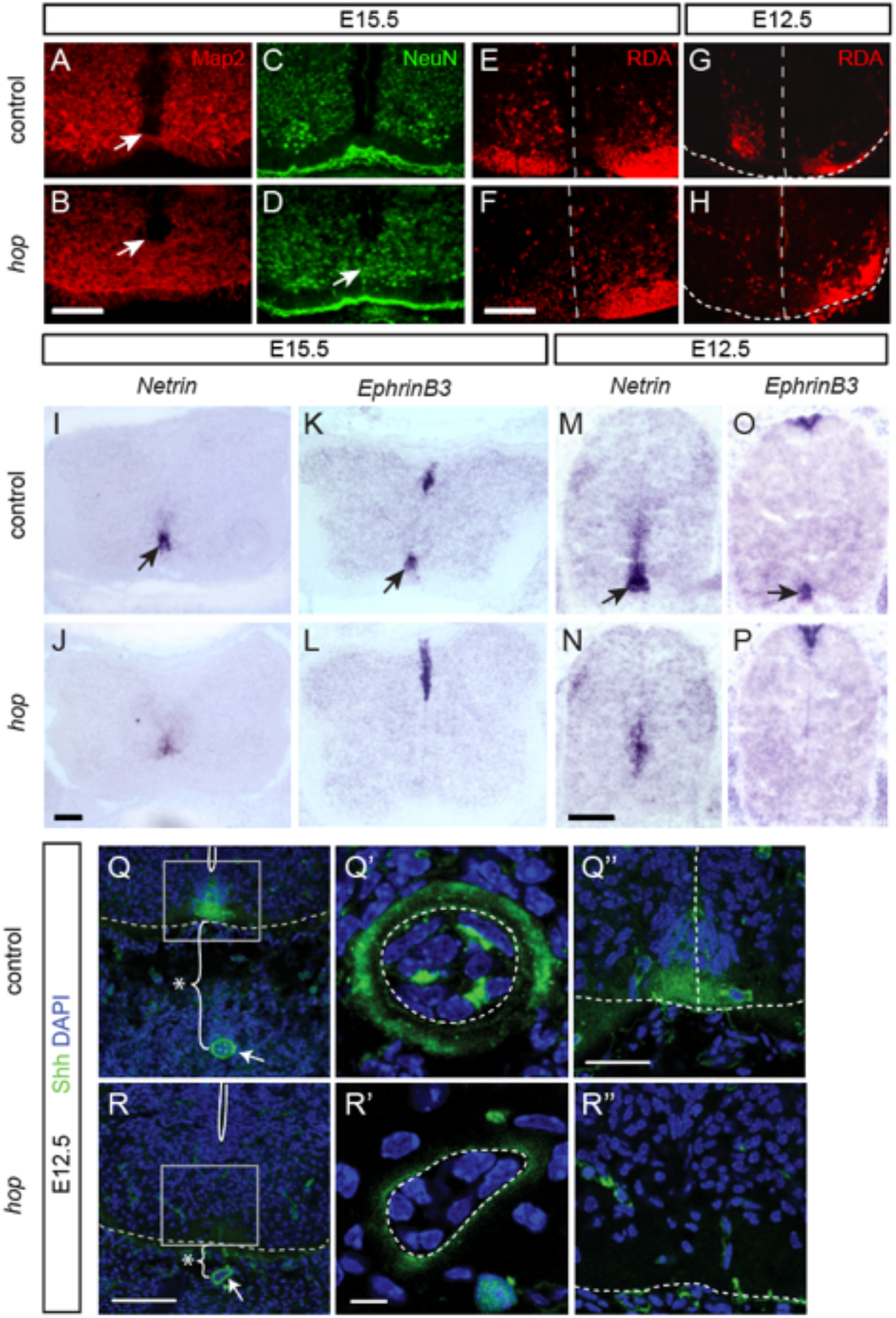
The distinct spinal cord phenotype in *hop* mice is present at embryonic stages E15.5 and E12.5. (A-D) Photomicrographs of immunohistochemistry staining on E15.5 *hop* and control spinal cords with Map2 (red) and NeuN (green) showing aberrant fibres and misplaced neurons not respecting the ventral midline. Arrows in (A, B) point to the central canal, arrow in (D) point to neurons on the midline. (E-H) RDA retrograde CIN tracing on E15.5 (E-F) and E12.5 (G-H) spinal cord. Dotted line indicates midline and outline of the spinal cord. (I-P) In-situ hybridization on E15.5 (I-L) and E12.5 (M-P) spinal cords with reduced Netrin1 and EphrinB3 levels in *hop* mice (J, L, N, P) compared to control (I, K, M, O). Arrows pointing to ventral Netrin-1 and Ephrin B3 mRNA expression (I, K, M, O). (Q,R) Photomicrographs of Shh immunohistochemistry staining on transverse spinal cord sections show notochord and floorplate phenotype, in low power (Q,R) and close up from control (Q’, Q’’) and *hop* (R’, R’’). Decreased distance between the notochord and spinal cord is indicated by the stars. Reduced amounts of Shh and malformed notochord in hop mice are pointed out by arrows and seen in the enlargement (Q’,R’). Lack of floorplate cells and Shh in the spinal cord in *hop* mice (R’’) compared to control (Q’’). Scale bars 50 μm (A-H), 100 μm (I-R), 25 μm (Q’,R’) and 30 μm (Q’’, R’’).

Several synchronous locomotor phenotypes in mice have been associated with mutations involving midline axon guidance molecules, including Netrin1 and ephrinB3. The absence of the midline chemo-attractant Netrin1 leads to a complete switch from alternating to synchronous fictive locomotor activity (10). Further, mutations in either the ligand ephrinB3 or the receptor EphA4 cause abnormal midline crossing of corticospinal axons and interneuron axons in the CPG, resulting in mice with a characteristic hopping gait phenotype (3). To assess the expression of axon guidance molecules in the midline of *hop* mice, we performed *in-situ* hybridization experiments on E15.5 and E12.5 spinal cord tissue with *Netrin1* and *EphrinB3* mRNA probes. The results showed that both *Netrin1* and *EphrinB3* expression levels were severely reduced in the ventral spinal cord of *hop* mice compared to controls, while the expression of *EphrinB3* in the dorsal spinal cord remained (Fig. 4I-P). These data suggest an early axon guidance defect in the ventral part of the spinal cord, which could at least partly explain the physiological phenotype in *hop* mice.

However, we could not exclude the possibility that the reduced axon guidance molecule expression in the ventral spinal cord might be the consequence of earlier developmental errors. Therefore, we continued exploring the expression of Shh, a key regulator of early development. Immunohistochemistry on E12.5 tissue with a Shh antibody revealed that Shh levels were reduced in both the notochord and spinal cord of *hop* mice (Fig. 4Q, R). Strikingly, in the ventral spinal cord of *hop* mice, Shh expression was absent, likely as a result of a missing floorplate (Fig. 4Q’’, R’’). Moreover, notochord morphology was abnormal displaying disorganised cells and low Shh expression (Fig. 4Q’, R’). Furthermore, the distance between the notochord and ventral spinal cord border was clearly decreased (Fig. 4Q, R).

### Identification and validation of the *hop* mutation

Our morphological characterization of the *hop* mouse pointed to a Shh dependent developmental origin for the observed phenotype. In parallel to our phenotypic analysis, we bred *hop* mice, which have an undefined mixed background but were obtained on balb/c, with a C57BL/6 mouse strain and generated 800 second generation offspring (F_2_) for phenotypic analysis. The pedigree was screened with single nucleotide markers from the proximal end of chromosome 6. This produced sufficient recombination events to map the *hop* mutation between 38.8 Mbp and 43.8 Mbp on chromosome 6, a 5 Mbp region containing 170 genes (Fig. S5). The genes in the candidate region were bioinformatically evaluated and categorized according to their functional annotation. *Ttc26*, starting at 38.4 Mbp, has been classified as an intraflagellar transport (IFT) complex B protein (48). Studies conducted on other IFT mutants have revealed phenotypic defects similar to those in the *hop* mutant including ventral spinal cord patterning defects, loss of ventral cell types and polydactyly (49–51). A point mutation in the *Ttc26* gene was found as in a previous study of *hop* mice (32). Our own whole exome sequencing of a *hop* mouse and of *Ttc26* exon 15 in several heterozygous (n=3) and homozygous (n=3) individuals confirmed a C to A transversion at position 38,412,065 bp causing a premature stop codon and a truncated form of the TTC26 protein (Fig. S5).

Thus, in the *hop* mouse, defects that are associated with early developmental patterning and Shh signalling are apparent, supporting the hypothesis that the identified point mutation in *Ttc26* is the causative mutation, and in line with a previous study (32). However, it is possible that other mutations are present in neighboring genes which may be adding to or even causing the observed phenotype. In the *hop* mutation area and in the close vicinity to the *Ttc26* gene, genes that have been associated with gait defects can be found. For example, *Hipk2* has been associated to TGF-beta signaling, another influential spinal cord morphant, and plays an essential role in the regulation of survival in the nervous system (52–54). Further, ablation of *Hipk2* generated mice with an ataxic-like phenotype implying that the *Hipk2* gene may contribute to the *hop* phenotype (55).

We sought to exclude such additional possible causative mutations and determine beyond doubt whether the underlying genetic cause in the *hop* mouse is the identified point mutation in *Ttc26*. Using clustered regularly interspaced short palindromic repeat/CRISPR-associated 9 (CRISPR/Cas9) technology, we introduced the identified point mutation within *Ttc26* exon 15 at position 38,412,065 bp in wild type mice (Fig 5A). This generated CRISPR *Ttc26*^*Y430X*^ mutant mice, which were smaller at birth and displayed polydactyly of all feet (Fig. 5L, M) highly reminiscent of *hop* mice, confirming that *Ttc26* point mutation is the causative mutation.

**Figure 5:**
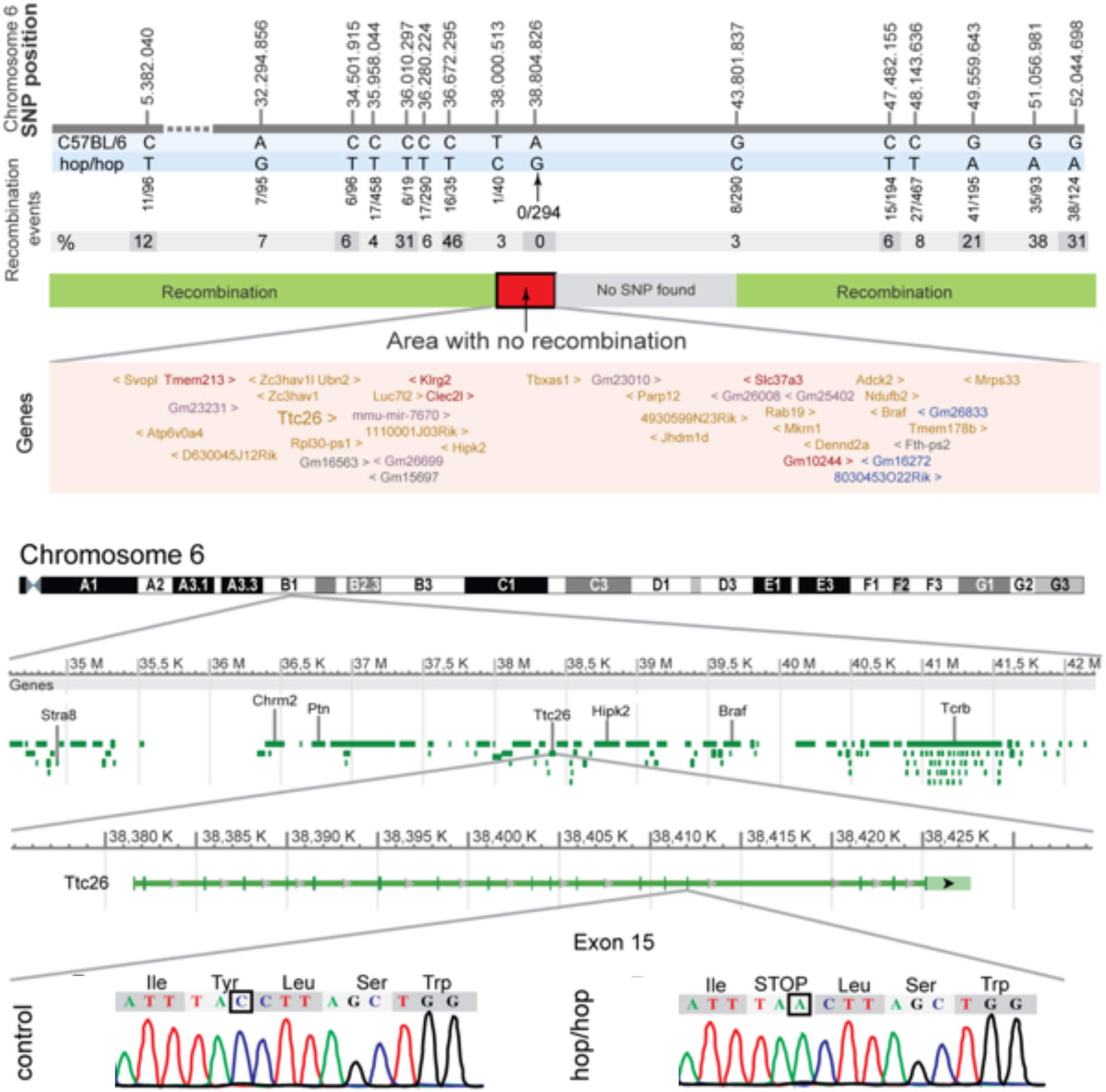
Introduction of a point mutation in *Ttc26* and analysis of the *hop* and *Ttc26*^*Y430X*^ mutants. (A) We applied the CRISPR/Cas9 system to generate mice with a point mutation in exon 15 of the *Ttc26* gene, generating a substitution of tyrosine to a premature stop codon. By simultaneous microinjection of guideRNA, hCas9 mRNA and a single-stranded donor oligonucleotide (ssDON) into mouse oocytes, mouse chromosome 6 harbouring the *Ttc26* gene was modified (left). The exon (black) and intron (red) organization is indicated together with Sanger sequencing traces of wild type (top) and the mutated allele (bottom) highlighting the premature stop mutation. The introduction of a PvuII site for screening purposes (blue) and the location of the guideRNA are indicated. (B-H) Immunostaining on embryonic spinal cord sections with *Ttc26*. Photomicrographs of immunohistochemistry experiments on E10.5 and E12.5 spinal cord transverse sections using antibodies as indicated. The expression of Nkx2.2 is absent in sections from *hop* mice at E10.5, while *Ttc26* is seen in the notochord both in control and *hop* sections. White box in (B,F) indicate areas of higher magnification in respective lower panels (B’-H’). For clarity, blue nuclear counterstain with DAPI has been omitted from the lower panels. Dashed lines outline the spinal cord and notochord. (I-K) Photomicrographs of Hematoxylin and Eosin staining on E16.5 transverse spinal cord sections. (I) Lumbar spinal cord of control mice showing a well delineated central canal (upper black arrow), a present ventral funiculus with a border between white and grey matter (dashed line) and a present ventral spinal artery (lower arrow). (J) Lumbar spinal cord of *Ttc26*^*Y430X*^ mice showing a poorly delineated dorsally shifted central canal (upper arrow), an absent ventral funiculus (upper star) and an absent ventral spinal artery (lower arrow). (K) Cervical spinal cord of *Ttc26*^*Y430X*^ mice showing a well delineated central canal (upper arrow) and a present ventral funiculus with a clear midline border (lower arrow). (L,M) Images of E12.5 (L) and E16.5 (M) *Ttc26*^*Y430X*^ mice showing preaxial polydactyly (arrowheads, star). (N,O) Photomicrographs of immunohistochemistry experiments on E16.5 transverse lumbar spinal cord sections of control and *Ttc26*^*Y430X*^ mice. (O) NeuN staining showing a poorly defined ventral midline in *Ttc26*^*Y430X*^ mice, with misplaced neurons within the white matter of the ventral funiculus. Enlargements on the right show cells positioned at the location of the missing midline in mutant but not control spinal cord sections. Scale bars (B-H, I-K, N,O) 100 μm, (B’-H’) 30 μm.

We next evaluated protein expression of TTC26 in the early developing spinal cord, together with markers for the ventral-most V3 progenitor domain, floorplate and notochord (Nkx.2.2 and Shh; Fig. 5B-H). Immunohistochemical analysis using antibodies against TTC26 showed that the truncated TTC26 protein was still detectable in *hop* mice (Fig. 5C’). Furthermore, E10.5 *hop* mutant embryos lacked the Nkx2.2 labelled progenitor domain when compared to control littermates, and the TTC26 immunopositive notochord was closer to the ventral floorplate in the *hop* mutant embryos compared to controls (Fig. 5B-C). Shh immunolabeling was found in the notochord and floorplate of control animals whereas it was entirely absent in *hop* mutant embryos (Fig. 5D, E). At E12.5, the improper positioning of the notochord and missing floorplate was further emphasized, as well as the detection of a misshaped notochord (Fig. 5F, G), however, these defects were not found in the thoracic region (Fig. 5H).

CRISPR *Ttc26*^*Y430X*^ mutant mice showed a spinal cord phenotype equal to the *hop* mice (Fig. 5I,J). Ventral spinal cord morphology matched the *hop* mice, with a poorly delineated and dorsally shifted central canal (Fig. 5I, J). Furthermore, like in *hop* mutants, defects were regionalized to the lumbar spinal cord with the cervical region unaffected (Fig. 5K). NeuN immunolabeling to identify neurons corroborated that the border between the white and grey matter was poorly defined, as was found in the *hop* mutants (Fig. 5N, O). There were also less myelinated structures and misplaced neurons within the white matter, resulting in an absent ventral funiculus (Fig. 5J, O). These findings demonstrate that the spinal cord phenotype found in *hop* mice was mirrored in *Ttc26*^*Y430X*^ mice. Thus, we conclude that the nonsense C-to-A point mutation in the *Ttc26*-gene, exon 15 is the sole cause of the observed spinal cord defects in *hop* mice.

Loss of Shh in mice results in degeneration of the notochord, absence of a morphologically distinct floorplate and abnormal dorsoventral patterning (39). Consequently, we analyzed embryonic spinal cords of *hop* mice using antibodies against transcription factors as markers for specific developmental subpopulations to assess patterning defects (Fig. 6, (56,57). We used Brn3a to examine the dorsal subpopulations dI1–dI3 and dI5 interneurons, which were largely unaffected, except in the ventral-most regions (Fig. 6C, D). Similarly, the dI4 – dI6 subpopulations labelled by Lbx1 (Fig. 6E, F) and the dI4, dI6, p0 and p4 subpopulations labelled by Pax2 (Fig. 6G, H) were present and appeared normal, except for the ventral-most labelled cells, which were found in close vicinity to the midline in *hop* mice. Markers for ventral V0v cells labelled by Evx1 and for dI3 and motor neurons labelled with Isl1/2 identified cells residing on and close to the ventral midline in *hop* mice (Fig. 6J, L), which was not seen in controls (Fig. 6I, K). Most striking, this analysis revealed that the floorplate and the ventral most Nkx2.2 positive progenitor domain were undetectable in *hop* mice, although present in controls (Fig. 6M, N). Thus, our results show that there is an early patterning defect in the lumbar spinal cord of *hop* mice, likely originating from a defect in the notochord early organizer.

**Figure 6:**
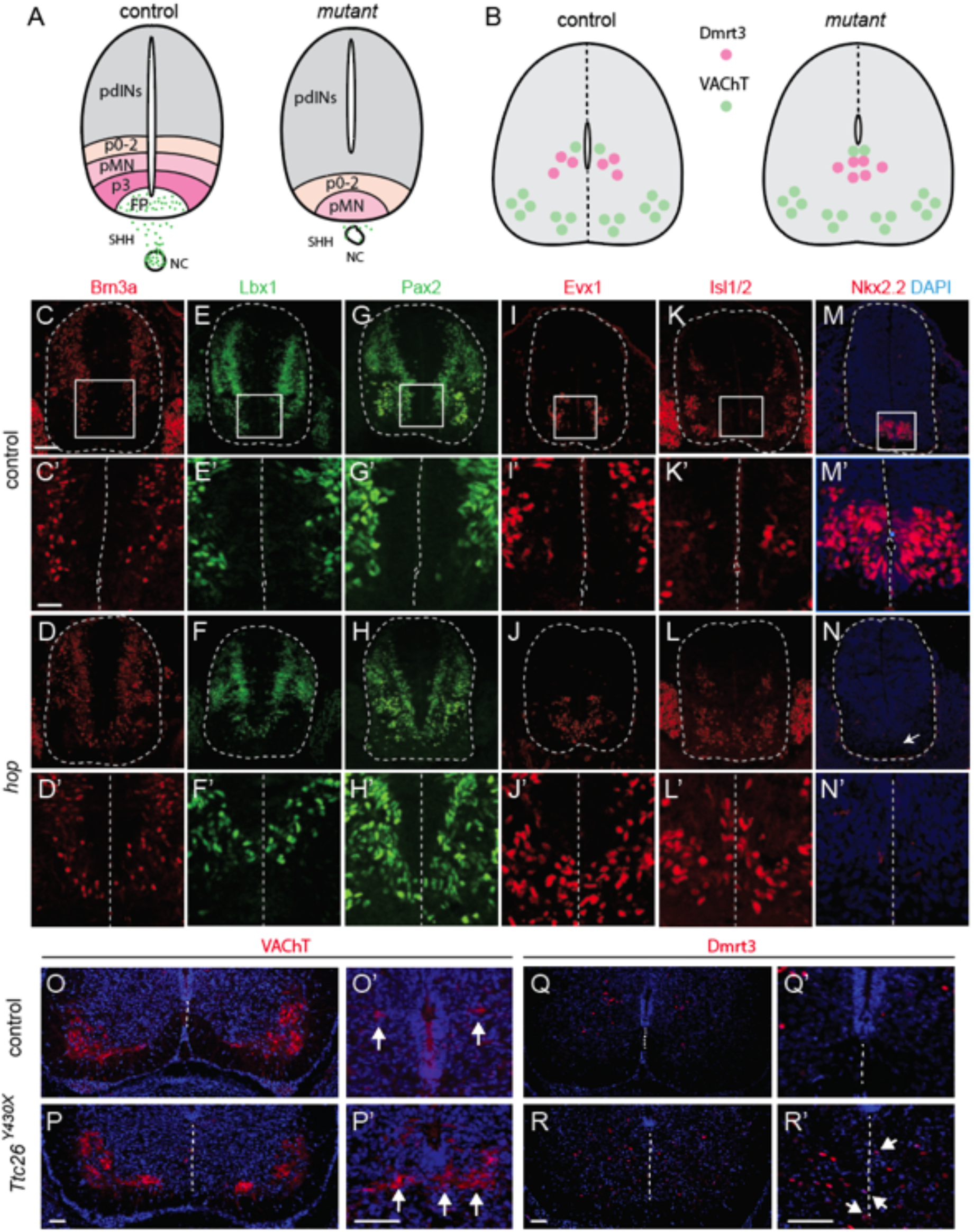
Ventral lumbar spinal cord phenotype in *hop* and *Ttc26*^*Y430X*^ mutant mice. (A,B) Schematic drawings of patterning and cell specification in the spinal cord at E12.5 (A) and E16.5 (B). (A) A gradient of SHH establishes positional information in the dorsoventral axis of the neural tube and establishes ventral progenitor domains for interneurons (p0, p1, p2, p3) and motor neurons (pMN). Progenitor domains and the subsequent neuronal subtypes arising can be recognized by their specific expression of transcription factors. Schematic summary of defects found in the *hop* mice (right) in the lumbar part of the spinal cord including weak Shh expression, misplaced and malformed notochord, a dorsally shifted central canal, absence of floorplate and p3 domains and absence of a ventral midline; pdINs, progenitor domains for dorsal interneurons, fp, floor plate. (B) Cholinergic neurons as identified by their VAChT expression (green) were found in the medial and lateral motor neuron cluster, as well as in neurons around the central canal. Dmrt3 expressing interneurons (pink) were found medioventrally from the central canal. In mutant *Ttc26*^*Y430*^ spinal cords (right), medial and lateral motor neuron nuclei were located at their expected positions. In contrast, central canal cholinergic neurons and Dmrt3 expressing interneurons were found ventral of the central canal and positioned in close proximity or even on top of the presumed position of the midline. (C-N) Immunostaining on E12.5 spinal cord sections revealed a missing Nkx2.2 progenitor domain and a migration defect of neurons in the lumbar spinal cord of *hop* mice. Photomicrographs of immunohistochemistry experiments on E12.5 spinal cord transverse sections using antibodies against transcription factors as indicated to the left. White boxes in (C-M) indicate areas of higher magnification in the respective panel below (C’-M’). Arrow in (N) points to missing Nkx2.2 progenitor domain. The stainings show misplaced cells and lack of ventral midline for the other tested progenitor markers. The outline of the spinal cord, central canal and midline are indicated by dashed lines (C-N’). (O-P) VAChT immunostaining showing no apparent differences regarding position and size of motor neurons between *Ttc26*^*Y430*^ and control mice. (O’,P’) Central canal VAChT expressing cells, as well as (Q-R) Dmrt3 expressing cells, were ventrally and medially displaced in *Ttc26*^*Y430*^ spinal cords compared to controls. (Q’,R’) Enlargement of the ventromedial area of the spinal cord demonstrating misplaced cells in *Ttc26*^*Y430*^ spinal cords (R’, arrows). Scale bars (C-N) 100 μm, (C’-D’) 50 μm and (E’-N’) 30 μm, (O) 300 μm, (O-R) 50 μm, (O’-R’) 100 μm.

Finally, we sought to investigate how two populations of direct importance for locomotion were affected: motor neurons and the dI6 inhibitory interneuron population (13,58). Immunohistochemistry using primary antibodies against vesicular acetylcholine transporter (VAChT) were used to label and count motor neurons in the lumbar part of *Ttc26*^*Y430X*^ and control spinal cords at E16.5. The count gave a mean value of 19.6 ± 9.4 motor neurons per hemi-section (51 hemisections counted) in the *Ttc26*^*Y430X*^ spinal cord and 20.6 ± 9.3 motor neurons per hemi-section (57 hemisections counted) in the control, which was not significantly different (independent-samples Student’s t-test; p=0.60). Examination of the motor neuron distribution in the lumbar region of the Ttc26^Y430X^ spinal cord revealed two distinct populations on each side of the midline, similar to controls. This indicates that, despite the midline fusion in the ventral spinal cord, the left and right populations of motor neurons remained separate, as was also found in the hop adult mutants (Fig. 3). We next explored the distribution of dI6 interneurons using anti-Dmrt3 antibodies. As expected, in controls they were located medioventrally in the spinal cord and formed two distinct populations on each side of the midline. However, in *Ttc26*^*Y430X*^ spinal cord, the Dmrt3^+^ interneurons were clustered on the midline were found further ventrally in the spinal cord (Fig. 6R).

## Discussion

Here we analyzed *hop* mice, focusing on the underlying cause for their aberrant locomotor phenotype. Both behavioral tests and fictive locomotion experiments showed that *hop* mice move using their hindlimbs in synchrony. Our developmental studies revealed an abnormal notochord, missing floorplate and Nkx2.2 progenitor domain in *hop* mice as well as a disorganized dorsoventral patterning with neurons residing on the ventral midline. Further, we found reduced expression of the axon guidance molecules netrin-1 and ventrally absent ephrinB3. By verifying the causative mutation in *Ttc26*, we revealed its important role for the formation of the lumbar locomotor network.

### Ventral spinal cord fusion explains the synchronous gait

According to the half-centre model of the spinal locomotor CPG proposed by Graham-Brown (1), the rhythmic pattern of alternating bursts of flexor and extensor muscles is produced by two neural populations in each half of the spinal cord. Commissural inhibitory and excitatory projections ensure normal left-right coordination (59). After cutting the ventral commissure the normal left-right circuit disappears indicating that ventromedially located CINs are indispensable for the bilateral coordination (12,18). Our morphological studies of the spinal cord revealed defects in the ventral spinal cord, including an absent midline and undefined white/grey matter border. The disturbed expression of axon guidance molecules in *hop* mice, seem to cause the observed aberrant fibres of short-, long- and locally-projecting INs. Further, their cell bodies were spread on the ventral midline compared to the two separate ventromedial clusters of commissural INs seen in control mice (Fig. 3, Fig. S4). Also, both excitatory and inhibitory neurons were found to reside on the ventral midline. This was further tested by adding drugs to increase inhibitory drive, which reverse synchrony back to alternation in other mutant mice with synchronous gait, demonstrating the presence of aberrant midline crossing of excitatory fibers (3). However, adding such drugs did not switch the synchronous pattern back to alternation in *hop* mice, ruling out a similar cause for their synchronous gait (Fig. 1).

Midline defects were discernable as early as E12.5, where we found that ventral neuronal progenitors in the lumbar spinal cord were located at the previous position of the missing midline (Fig. 6). This was also manifested four days later when the Dmrt3 interneurons originating from the dI6 progenitor domain assembled close or on top of the missing midline. These results indicate that in *hop* mice, the neurons that form the two separate CPG half-centres might functionally share part of the neurons, most likely solely caused by intermingling of left and right interneuron populations. This would essentially cause a functionally fused CPG center, with a synchronous output and gait as a consequence.

Expression of pre-pattern genes, along the dorsoventral axis of the spinal cord, is controlled by long ranging signals from the opposite poles of the neural tube. *Sonic Hedgehog, Gli3* and *Smoothened* are among the most important molecules for ventral cell differentiation, with the most ventral progenitor domain requiring the highest concentration of *Shh* to develop normally (60,61). Already at E10.5, the most ventral Nkx2.2 progenitor domain, giving rise to V3 neurons, was absent in the *hop* mice. V3 neurons are one of the major classes of excitatory commissural neurons in the mouse spinal cord and are components of the locomotor commissural network, with a role in establishing a stable and balanced locomotor rhythm (9,62).

Motor neuron populations, however, were not assembled at the midline in the mutant mice. Previous studies have shown that the upper edge of the Islet-1^+^ expressing motor column is dorsally shifted in Netrin-1 mutant spinal cords at E10 (63). In *hop* mice, at E10.5 and E12.5, cells expressing Islet-1^+^ were dispersed dorsally and clustered together on the ventral midline (Fig. 6K-L). Likewise, Robo mutant embryos (E9.5-E10.5) have a pack of misplaced Islet-1^+^ motor neurons within the floorplate. However, the ectopic Islet-1^+^ cells of Robo mutants are no longer visible in the floorplate by E12.5. This implies that the cells either die, turning off the Islet1 transcription factor, or migrate out of the floor plate between E10.5 and E12.5 (63). In accordance with these data, previously reported analyses of Slit/Robo mutant mice (E12) have failed to observe the clustering of motor neurons on the midline (64). Our study of the *Ttc26*^*Y430X*^ spinal cords at E16.5 suggest that decreased levels of Netrin-1 in fact do affect the initial positioning of motor neurons, but that compensatory mechanisms then correct the mis-placing before birth. We did not find a significant difference between the number of motor neurons in *Ttc26*^*Y430X*^ and control spinal cords, which indicates that the misplaced neurons most likely migrate back to their correct location, rather than undergo apoptosis. In addition to Semaphorin and Slit signalling governed by the transcription factors Islet1 and Islet2 (65,66), motor neuron positioning is regulated by Reelin and Cadherin signalling causing a secondary reorganization of motor columns into sub-clusters consisting of motor pools (67–69). The migration and positioning of motor neurons seems to be more strictly regulated than the positioning of CINs, possibly explaining why motor neurons are not clustered on the ventral midline of *Ttc26*^*Y428X*^ spinal cords. Further studies are needed to understand the exact underlying mechanisms for the observed differences between motor neuron and interneuron development in the ventral spinal cord.

### Rostrocaudal specificity of the midline fusion

In addition to the previously reported characteristic hopping gait in *hop* mice (28,33), our gait analysis showed that adult *hop* mice alternated their forelimbs while they moved their hind limbs in synchrony. Further, we found that the pattern of flexor-extensor alternation during fictive locomotion was maintained following hemisection, suggesting that the circuitry coordinating ipsilateral alternation is intact in *hop* mutants (Fig. S2). These findings suggest that the defect behind the characteristic hopping phenotype is limited to the lumbar spinal cord controlling the left-right alternation of the hind limbs. This is in accordance with the lumbar specific defects observed in the *Ttc26* mutant mice early in development.

What might be the possible explanation for the rostro-caudal specificity of the defect? Primary cilia and Shh signalling are critical for proper patterning of the neural tube along the dorsoventral axis (70,71), and TTC26 is required for accumulation of Gli at the ciliary tip (32). Shh KO mice do not generate motor neurons, however, in Shh and Gli3 double KO mice there is a rescue effect where motor neurons are generated predominantly in the lumbar region (72). This substantiates the involvement of the Shh signaling pathway in differential development along the rostral−caudal axis, possibly due to an unknown caudal factor such as retinoic acid (72). An alternative explanation from work in vertebrates and flies suggests that changes in rostrocaudal positional information provided by *Hox* genes can impact locomotor behaviors through modifying CPG organization (73). Of note, the *Hox1* gene cluster is relatively close to *Ttc26* and possibly, disturbance of such regulatory elements may cause a rostro-caudal restriction of the phenotype.

### The mutation in *Ttc26* is causative for the spinal cord fusion

Shh is crucial for induction of both floorplate and dorsoventral patterning of the spinal cord (39,60,74), and in particular, defective secretion of Shh can result in the spinal cord developing without floorplate cells and V3 interneurons (75). *Shh* mutant mice have several defects in midline structures such as the notochord and cyclopia (76). A similar spinal cord phenotype is present in HNF-3β null mutant mice. HNF-3β is a transcription factor expressed in the three midline organizing centers; the node, notochord and floorplate (77). Similar to Swidersky et al., 2014, our positional cloning analysis localized the hop mutation to *Ttc26*. TTC26, a component of Intraflagellar Transport (IFT) complex B, is a structure necessary for cilia formation and protein transport, and is important for Shh signaling. In addition, Ttc26 (also named IFT56) regulates vertebrate developmental patterning by maintaining IFTB complex integrity and ciliary microtubule architecture (78). Moreover, cilia motility is important for embryonic left-right determination (79). Although it is likely that the found mutation is the causative one, we noticed some differences between our analysis and the Swiderski et al. (2014) study. The spinal cord ventralisation was found in both studies, but additionally, we found midline fusion defects as well as lack of the ventral midline and Nkx2.2 positive V3 progenitors in the lumbar spinal cord. These differences opened the possibility of other additional mutations in the *hop* mice. For example, *Gli1*^*−/−*^; *Gli2*^*+/−*^ mice have a hopping gait (80), and *Gli3*^*+/−*^ mice have preaxial polydactyly (80,81). In addition, a similar distortion of notochord, midline fusion (E14) and missing Nkx2.2 domain has been observed in *Gli1/2* double mutants. Also, in *Gli2*^*-/-*^ mouse embryos, the most ventral cells (the floor plate) are not specified (80).

The lower expression levels of Shh seen in *hop* mice could also be caused by mutations in upstream regulators of Shh, events that could take place earlier in development. As mentioned above, the HOX1 cluster, located on cytoband B3-C >25.40 cM in the vicinity of the *hop* region, could be affected in the *hop mouse* mutation. Mice with homozygous mutations in one or multiple members of the cluster display a range of phenotypes including developmental defects in the skeletal, reproductive and nervous systems (82). Two other genes in the *hop* region are associated to gait defects or could explain a fused spinal cord midline. When the *Braf* gene is mutated it has been reported to cause developmental deficits such as craniofacial dysmorphism and extra digits (83). Mice with a targeted mutation in the transcriptional cofactor homeodomain interacting protein kinase 2 (HIPK2) display abnormal gaits including shuffling, reduced width and short stride length (54). However, by CRISPR technology we introduced a point mutation in the *Ttc26* gene, which produced a premature stop codon mimicking the *hop* mouse mutation. Such mice displayed the same spinal cord ventral midline fusion as found in *hop* mice and thereby established that such a point mutation sufficiently explains the mutant phenotype.

### Concluding paragraph

To our knowledge, the *hop* mutant is the first model with a specific ventral spinal cord fusion, in which the components of the CPG can be studied and understood. Our data support the two half-centre hypothesis by demonstrating that midline separation is crucial for left-right alternation of locomotion. Despite the loss of the ventral midline, functional neuronal networks develop and form connections able to produce coordinated, albeit synchronous, activity. These studies together with our findings indicate that the aberrant spinal cord phenotype observed in *hop* mice is likely the result of abnormal developmental processes including induction from the notochord and Shh signalling.

## Material and Methods

### Animals

#### Mice

CByJ.Cg-hop/J mice were imported from Jackson Laboratory (N12 on a backcross-intercross to BALB/cByJ). The colony was kept on the BALB/cByJ background for analysis of the locomotor phenotype and outcrossed to the C57BL/6J background for positional cloning. *Ttc26*^*Y430X*^ were produced by Siu-Pok Yee at the Gene Targeting and Transgenic Facility, Uconn Health Center. All experiments involving animals were approved by the appropriate local Swedish ethical committee (C147/7 or C79/9), and by the University of Calgary Health Sciences Animal Care Committee.

### Gait study

2-3 month old animals were trained to walk on a 1 meter long and 10 cm wide track the day before the experiment. Mice were taken up and painting of the paws was mimicked with a brush. Each mouse had to go to the end of the track at least 3 times without stopping. The day of the experiment mice were handled exactly the same as during the training but this time the paws were painted with two distinct colors for the forelimbs (red) and hindlimbs (blue). Footprints were recorded on Whatmann paper placed at the bottom of the track. For each animal, 6 runs were recorded and analyzed according to (Kullander et al., 2001).

### Electrophysiology

Experiments were performed on 0-3 days old (P0-P3, weight 1.16 g to 4.05 g; n = 62) *hop* homozygote, heterozygote and wild type control mice. Mutant mice were identified by preaxial polydactyly of the hindfeet (Fig. S1) and the synchronous activity in their hindlimbs when suspended in the air with their tails lightly pinched. There were no obvious deficits in gait in the *hop* heterozygote and wild type control mice.

For in vivo experiments, animals were anaesthetized by hypothermia and then suspended in a sling such that the fore- and hindlimbs moved freely (84). Electromyographic (EMG) electrodes were inserted into the left and right tibialis anterior muscles parallel to the muscle fibres, and a grounding electrode was inserted subcutaneously into the back. EMG electrodes were made of 75 mm Teflon-coated platinum-iridium wires (A-M Systems Inc.). A heat lamp was used to keep the air temperature at 30°C. Recordings were amplified (1000 times), bandpass filtered (100 – 1 kHz), and digitized at 3 kHz (Axon Digidata 1322A) for future analysis. 5 – 10 minutes after the EMG wires were inserted, air-stepping sequences were elicited by pinching the tail with forceps and the activity was recorded.

For in vitro experiments, the animals were anaesthetized by hypothermia. Animals were rapidly decapitated, eviscerated and the remaining tissue was placed in a dissection chamber filled with oxygenated (95% O_2_ – 5% CO_2_) artificial cerebrospinal fluid (ACSF: concentrations in mM: 128 NaCl, 4 KCl, 0.1 CaCl_2_, 2 MgSO_4_, 0.5 Na_2_HPO_4,_ 21 NaHCO_3_, 30 _D_-glucose). A ventral laminectomy exposed the cord, and the ventral and dorsal roots were cut. The spinal cord was transected at thoracic 1 – 3 (T_1-3_) to sacral 2 – 3 (S_2-3_) and carefully removed from the vertebral column. After 30 minutes, the preparation was transferred to the recording chamber and superfused with oxygenated ACSF (concentrations in mM: 128 NaCl, 4 KCl, 1.5 CaCl_2_, 1 MgSO_4_, 0.5 Na_2_HPO_4_, 21 NaHCO_3_, 30 _D_-glucose). The bath solution was then heated gradually from room temperature to 27°C. The preparation was allowed to acclimate for one hour thereafter. Population motoneuron bursting activity was recorded using suction electrodes into which segmental ventral roots from the left and right lumbar L_2_ and L_5_ segments were drawn (34). The resultant neurograms were amplified (100 – 20,000 times), band pass filtered (100 Hz – 1 kHz) and digitized at 2 – 5 kHz (Axon Digidata 1320) for future analysis. Ten minutes of control baseline activity was recorded prior to adding drugs. 5 mM N-methyl-_DL_-aspartic acid (NMA, Sigma-Aldrich), 10 – 20 mM Serotonin (5-HT, Sigma-Aldrich) and 50 – 75 mM dopamine (DA, Sigma-Aldrich) were added to the bath and the rhythm stabilized for 10 – 30 minutes. In some experiments, a GABA uptake inhibitor (0.1 – 1 mM nipecotic acid or 50 – 100 mM NO-711, Sigma-Aldrich) and/or a glycine uptake inhibitor (100 mM sarcosine, Sigma-Aldrich) was added to the bath solution. In some experiments, we examined whether a hemisected spinal cord was capable of generating coordinated rhythmic activity. In these experiments, we first examined the activity patterns in the presence of rhythmogenic drugs. We then removed the recording electrodes and using a pair of micro-clippers completely mid-sagittally hemisected the spinal cord. The suction electrodes were then reattached to the ventral roots.

Data were digitally rectified, integrated and then analyzed using custom written programs (MatLab, MathWorks, Natick, MA). Locomotor-like activity was quantified using time series analysis. Time series analysis was performed by taking intervals of 60 seconds of raw data, rectifying the data, applying a low-pass filter and resampling at 100 Hz. Means were subtracted from the processed data and further smoothed using a digital filter (Savitzky-Golay, 3^rd^ order polynomial operating over 13 points). Cross and auto-correlograms were then calculated and the quality of the rhythm was assessed by measuring the correlation coefficients for the segmental L_2_ ventral root bursts and the left L_2_ and left L_5_ ventral root bursts. To measure rhythm stability, the peak-to-trough correlation coefficient (PTCC) was calculated from the cross correlogram by subtracting the minimum negative value of the correlation coefficient from the maximum of the first positive peak over the first 75 lags (each lag = 50 ms). Stable synchronous rhythms typically have high positive correlation coefficients at zero phase lag. The cycle periods for the resultant rhythm were calculated by measuring the number of lags from one peak to the next from the auto-correlogram. The phase lag between ventral root bursts was obtained from the cross correlogram and was defined as the distance from the minimum trough around lag 0 to the next peak divided by the cycle period (85). Data are expressed as mean ± standard error of the mean and significance was analyzed using paired and unpaired Student’s t-tests if the data were normally distributed (P<0.05). Data that were not normally distributed were analyzed using a Wilcoxon-Signed Rank test. Multiple data points were analyzed using a one way ANOVA followed by a Tukey posthoc test to detect significant differences. To analyze *in vivo* EMG data and to illustrate phase relationships for selected *in vitro* experiments, we used circular statistics, in which the phase was normalized from 0 to 1 (86). If the length of the arrow is large this suggests a tendency for the rhythms represented by the two neurograms to be coupled. Significance was computed using Rayleigh’s test (P<0.05).

### Tracing of commissural interneurons

Fluorescent dextran-amines 3,000 MW rhodamine-dextran-amine (RDA) and 3,000 MW fluorescein-dextran-amine (FDA) (Invitrogen, Sweden) were used for retrograde tracing of commissural interneurons (CINs) as described previously (87). P0–P3 mice spinal cords were prepared as described earlier (25). Tracings on E12.5 embryos were performed essentially the same way, but the spinal cord remained within the vertebral column during tracer application and incubation. Preparations were incubated for 12–16 h and then fixed in 4% paraformaldehyde (PFA) in 0.1 M phosphate-buffered saline (PBS), pH 7.4 and stored dark at 4°C for 1 week. Spinal cords were cut into 60 mm thick transverse sections on a vibratome (Leica, Germany) and stored in the dark at –20°C until analysis. Embryonic tracings were fixed for 2 hours, transferred to 30% sucrose in PBS at 4°C over-night, embedded in OCT and 12 mm sections were cut using a cryostat (CM1800, Leica), collected onto Superfrost slides (Menzel-Gläser, Germany)

### *In situ* hybridization

Free floating vibratome sections were rehydrated in consecutive washes for 10 min in 75%, 50% and 25% methanol in PBT, bleached in 6% hydrogen peroxide in PBT for 15 min and treated with 0.5% Triton X-100 for 5 min. The sections were digested with proteinase K (10 μg/ml) in PBT for 15 min. The digestion was stopped with a wash in glycine (Scharlau Chimie, Spain; 2mg/ml) in PBT for 5 min, and sections were postfixed in 4% formaldehyde for 20 min. The sections were prehybridized at 65°C in hybridization buffer (50% formamide, 5xSSC pH 4.5, 1% SDS, 50 μg/ml tRNA (Sigma) and 50 μg/ml heparin (Sigma)) for two hours prior to addition of probe. 1 μg/ml probe was added to the hybridization buffer and sections were hybridized over-night at 65°C. Excess probe was removed by washes in wash buffers (50% formamide, 5xSSC pH 4.5 and 1% SDS; 50% formamide, 2xSSC pH 4.5 and 0.1% tween-20) at 65°C for 3 times 30 min each. The sections were transferred to blocking solution (1% blocking reagent in TBST) for 2 h before addition of anti-DIG AP (1:5000) diluted in blocking solution and incubated over-night at 4°C. The sections were treated with levamisole (0.5 mg/ml) in TBST and levamisole (0.5mg/ml) in NTMT (100 mM NaCl, 10 mM Tris-HCl pH 9.5, 50 mM MgCl_2_ and 0.1% Tween-20) before developing in BM purple AP substrate (Roche) at 37°C 1h – 4 days. Between additions of new chemicals, sections were washed with either PBT (prior to addition of probe) or TBST (after addition of probe). In *situ* hybridization on cryosections was performed essentially as described (88). The VIAAT probe covers nucleotides 588-2072, the Vglut2 probe 1616 –2203 used as described earlier (89), the VAChT probe 1534 –2413, Netrin-1 and EphrinB3 probe 1-1021.

### Immunohistochemistry

The mouse tissue was cryoprotected in 30% sucrose in PBS, embedded in OCT medium and sectioned at a thickness of 12-14 µm on a cryostat (CM1800, Leica). The sections were washed in PBS and pre-blocked in blocking solution - 5% goat serum, 0.3% Bovine Serum Albumin (BSA, Sigma Aldrich) and 0.1% Triton^®^X-100 (Sigma Aldrich) in PBS. Primary antibodies were diluted in blocking solution, added to the sections and incubated over night at 4°C. The following dilutions were used: mNkx2.2 1:100 (Hybridoma bank), mEvx1 1:50 (Hybridoma bank), gpLbx1 1:10000 (kind gift from C. Birchmeyer), rPax2 1:1000 (Covance), mBrn3a 1:500 (Chemicon), mIsl1/2 1:100 (Hybridoma bank), mLhx1/5 1:500 (Hybridoma bank), Lhx2/9 1:8000 (kind gift from Dr. T.Jessell), mShh 1:100 (Hybridoma bank), mMap2 1:500 (Chemicon, Sweden), rbNeuN 1:1000 (Chemicon), MBP 1:500 (AbCam), rbTtc26 1:1000 (Novus Biologicals), mNeuN 1:500 (Chemicon), gpVAChT 1:500 (Millipore), gChAT 1:250 (Millipore), mParvalbumin 1:1000 (Sigma) and gpDmrt3 1:10 000 (custom made using the immunizing peptide CKQSIYTEDDYDERS-amide).

For some antibodies, antigen retrieval was necessary in order to get good immunohistochemical staining. The sections were then washed with PBS and thereafter placed in a metal rack inside a beaker filled with preheated citrate buffer (9 ml 0.1 M citric acid, 41 ml 0.1 M trisodium citrate dehydrate and 450 ml dH_2_O – pH6.0). After 30 minutes at 98°C, the beaker was left to cool down at room temperature for 30 minutes. Subsequently the sections were washed with dH2O for 1 minute and pre-blocked in 1% normal goat serum and 0.1% Triton^®^X-100 in PBS for 1 hour. Finally, the primary antibodies were diluted in blocking solution (1% normal goat serum and 0.1 % Triton^®^X-100 in PBS), added to the sections and incubated over night at 4°C. The following day, the sections were washed with PBS and incubated with DAPI (Sigma Aldrich, 200ng/ml) and the secondary antibodies (diluted 1:500 in blocking solution) for 2-3 hours at room temperature. After washing with PBS, the sections were mounted in Mowiol^®^ 4-88 (Sigma Aldrich). Secondary antibodies used: goat anti-mouse conjugated with FITC 488 (F(ab’)_2_Fragment, 111-096-003, Jackson Immunoresearch Labs), donkey anti-guinea pig conjugated with Alexa 594 (Jackson Immunoresearch Labs), donkey anti-rabbit conjugated with Alexa 488 (Invitrogen) and donkey anti-goat conjugated with Alexa 488 (Jackson Immunoresearch Labs). *Ttc26*^*Y430X*^ mice embryos (E16.5) were euthanized, genotyped, fixed in 4% paraformaldehyde for 4 hours on a rocking table at room temperature and then washed in 0.1 M phosphate buffered saline (PBS - Sigma Aldrich, Stockholm, Sweden) for 30 minutes before being cryoprotected in 30% sucrose in PBS. The tissue was shipped in 30% sucrose in PBS (in a falcon tube placed inside a Styrofoam box with cool-packs) to Uppsala University. Upon arrival 5 days later the embryos were frozen in OCT medium (Richard Allan Scientific NEG50™) on dry ice. Cryostat sections were cut (cryostat CM1800, Leica) at a thickness of 14µm and placed on Superfrost slides (Menzel-Gläser, Germany).

### Histology staining

#### Haematoxylin and Eosin staining

Cryosections of mouse spinal cord were rehydrated in dH_2_O for 1 minute, stained with progressive haematoxylin (Mayer’s) for 90 seconds respectively 6 minutes, washed twice in dH_2_O for 1 minute, dipped in Scott’s tap water (20 g MgSO_4_·7H2O and 2 g NaHCO_3_ in 1 L tap water), dehydrated in 96% Ethanol in dH2O for 1 minute and finally counter-stained with 0.5% Eosin (Sigma Aldrich) in dH_2_O for 20 seconds. Afterwards the sections were dehydrated with successive washes in 70% and 96 % Ethanol in dH2O for 1 minute and treated with X-tra Solve^®^(MEDITE) for 4 minutes before being mounted in Pertex® (Histolab®).

#### Other staining

3,3’-diaminobenzidinetetrahydrochloride (DAB) (Sigma, St Louis, MO, USA) was used on free-floating spinal cord sections to visualize white matter (dark brown) and Luxol fast blue (LFB) was used to analyze paraffin embedded brain sections staining myelin and phospholipids (blue-green) counterstained with cresyl violet as previously described (2).

### SNP Genotyping

A three-generation pedigree was generated after crossing one homozygous hop female with a C56BL/6 male. Four males and 8 females from the F1 generation were intercrossed. 780 F2 animals were genotyped for the selected SNPs by using a SNP Reagent Kit protocol (Pyrosequencing AB, Uppsala, Sweden) and custom TaqMan assay chemistry (Applied Biosystems).

### Exome sequencing of hop

Genomic DNA from a *hop* homozygote (stock 002718) and BALB/cByJ control (stock 001026) was obtained from the Jackson Laboratory’s DNA Resource. Exomes were captured at the Yale Center for Genome Analysis, using the NimbleGen SeqCap EZ Mouse Exome and following NimbleGen protocols. Captured pools were sequenced (75bp, paired-end) on an Illumina HiSeq 2000 using previously described methods (90). We obtained ∼91 million (BALB/cByJ) to ∼113 million (*hop*) high-quality reads. Illumina reads were first trimmed based on their quality scores to remove low-quality regions using the program Btrim (91). A cutoff of 20 for average quality scores within a moving window of size 5-bp was used. Minimum acceptable read length was 25-bp. Other parameters of Btrim were set to defaults. The pre-processed reads were then aligned to the mouse genome reference sequence (mm9) using the BWA mapping program (92). The mapping results were converted into SAMtools pileup format using SAMtools programs (93). PCR duplicates were removed using the rmdup command from SAMtools, resulting in ∼84x (BALB/cByJ) or ∼100x (*hop*) coverage across the exome. >97% of all bases included in the exome showed at least 8x coverage and >90% of the bases showed at least 20x coverage. Single nucleotide variations (SNVs) were called using SAMtools’ pileup command. Further filtering was performed using in-house scripts to exclude those SNV calls that had less than 3 reads or a SNP score less than 20. Annotation was added based on the UCSC RefSeq gene model (http://genome.ucsc.edu/ (94)). Based on SNV homozygosity mapping, the interval was narrowed to ∼700 Kb on Chromosome 6, (37.9-38.6Mb) which fell within the previously mapped *hop* genomic region. After filtering for known SNPs and repeat regions, we identified 17 possible point mutations, 16 of which were located in introns or 3’ untranslated regions. The remaining homozygous point mutation was located at Chr6:38,362,071 (**C>A**) and results in a nonsense mutation (**Y430X**) within the *Tetratricopeptide repeat protein 26* (*Ttc26*) gene.

### CRISPR mutagenesis

We used CRISPR/Cas9-mediated genome editing technology with the help from Siu-Pok Yee, Ph.D. at the Gene Targeting and Transgenic Facility, Uconn Health Center. A C/A transversion at position 38412065 in exon 15 of the gene *Ttc26* on chromosome 6 was created (95) to replace a TAC codon in the amino acid chain for tyrosine 430 with a TAA to create a premature STOP codon. A silent point mutation was also introduced to create a novel PvuII site, 3’ of the knock-in termination codon in *Ttc26* to allow for straightforward genotyping of the mice by PCR followed by a PvuII digestion. The PvuII site also affected the PAM sequence of the CRISPR so re-digestion of the KI allele was avoided. CAS nuclease, a guide RNA and an oligonucleotide carrying the desired mutation and homology arms were co-injected into the nuclei of fertilized oocytes, followed by implantation of the eggs into surrogate mothers to obtain offspring. An allele-specific primer was developed for genotyping of the pups. We obtained one *Ttc26* Y430X founder female that appeared to be heterozygous for the mutation (1:1 peak ratio from sequencing, Fig 5A). From F1 pups from the founder female, we bred heterozygous males and females to obtain homozygous *Ttc26*^*Y430X*^ mouse mutants.

### Imaging and picture processing

Fluorescent and bright field images were acquired either on an Olympus BX61WI microscope (Olympus, Sweden) using the Volocity software (Improvision, Lexington, USA) or on a MZ16F dissection microscope with a DFC300FX camera and FireCam software (Leica). Close-ups were taken using the OptiGrid Grid Scan Confocal Unit (Qioptiq, Rochester, USA) or a confocal microscope (Zeiss LSM 510 Meta, Oberkochen, Germany). Captured images were auto-leveled using Adobe Photoshop software.

## Acknowledgements

We thank C-J. Rubin, L. Strömstedt, A. Reis and S.P. Yee for technical assistance. We are also grateful to S. Weatherbee for discussions of earlier versions of this work.

## Supplementary material

**Figure S1:**
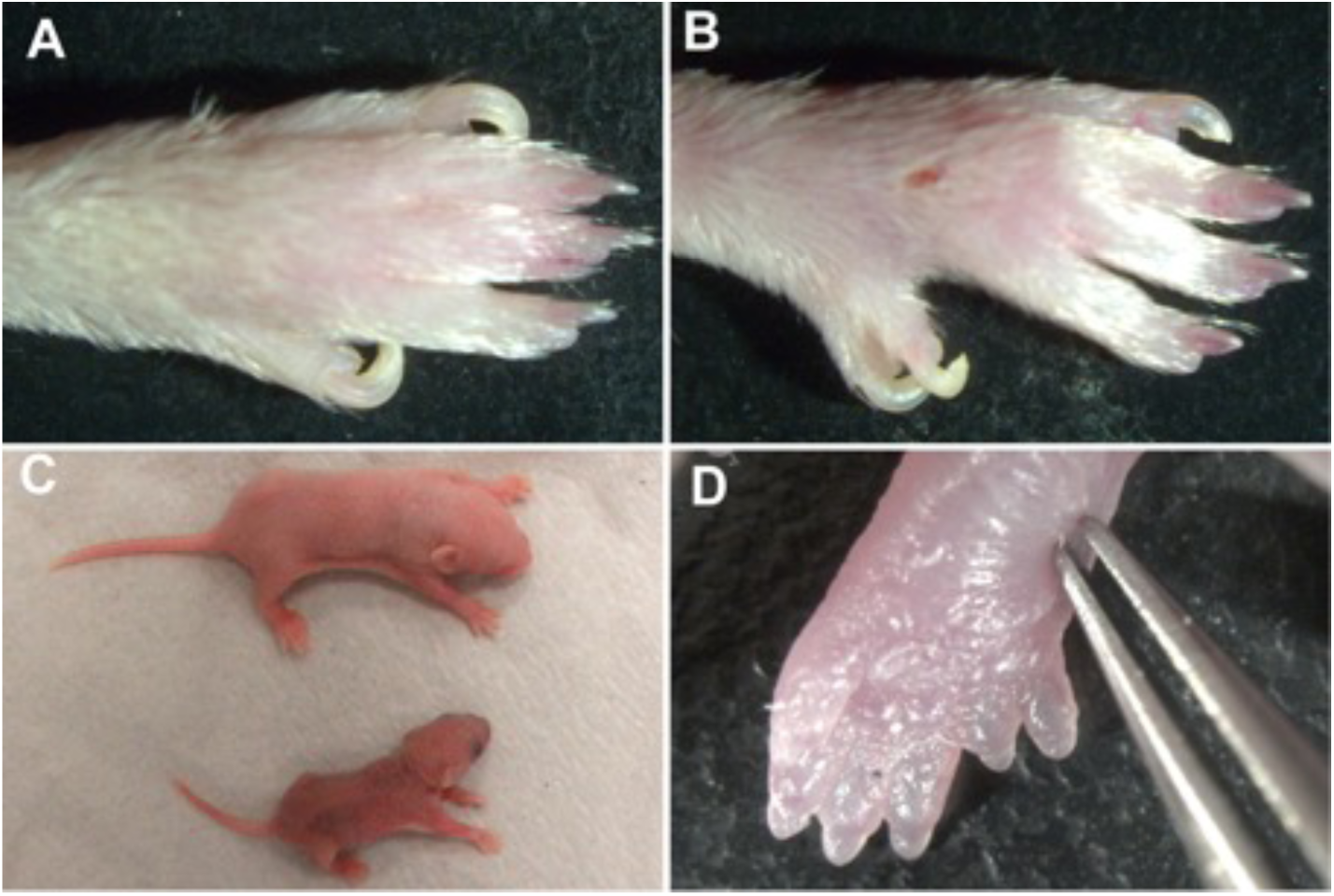
Gross *hop* mice phenotype. Photos of paws from a wild type (A) and *hop* (B,D) mice. *Hop* display preaxial polydactyly of all feet seen in the adult (B) and visible already at E15.5 (D). Comparison of size of newborn wild type (top) and *hop* mice demonstrate smaller size at birth (C).

**Figure S2:**
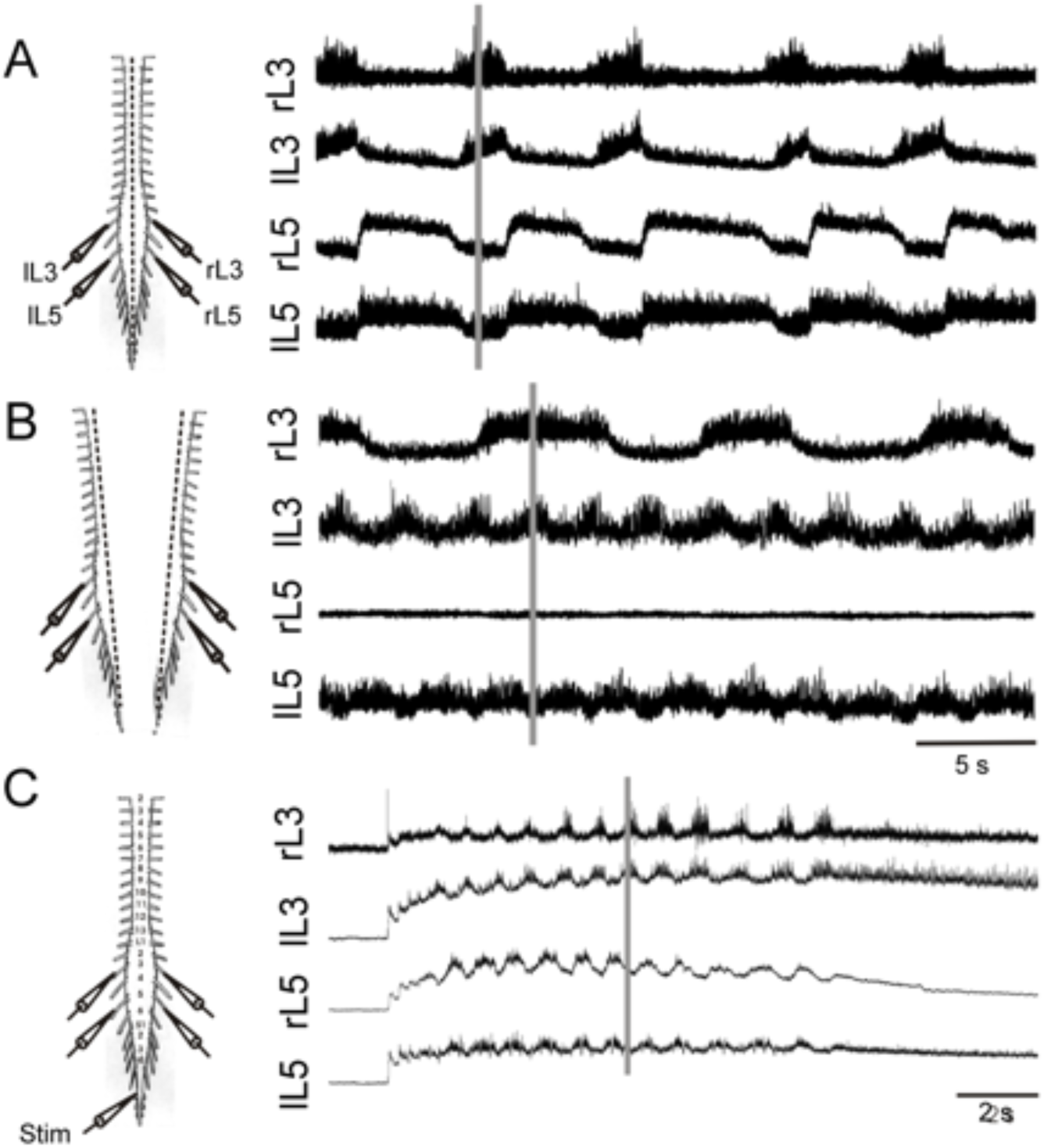
Mid-sagittal hemisection does not disrupt the ipsilateral alternating burst pattern. (A) Neurograms illustrating a typical pattern recorded from a *hop* mouse. (B) Pattern recorded from the preparation following a midsagittal hemisection. Note the preservation of the L3-L5 pattern suggesting that flexor-extensor coordination is maintained. (C) Schematic illustrating stimulation and recording arrangement for cauda equina stimulation. Cauda equina evoked rhythmic pattern (4 Hz, 10 s train) showed segmental synchronous pattern and ipsilateral alternating pattern.

**Figure S3:**
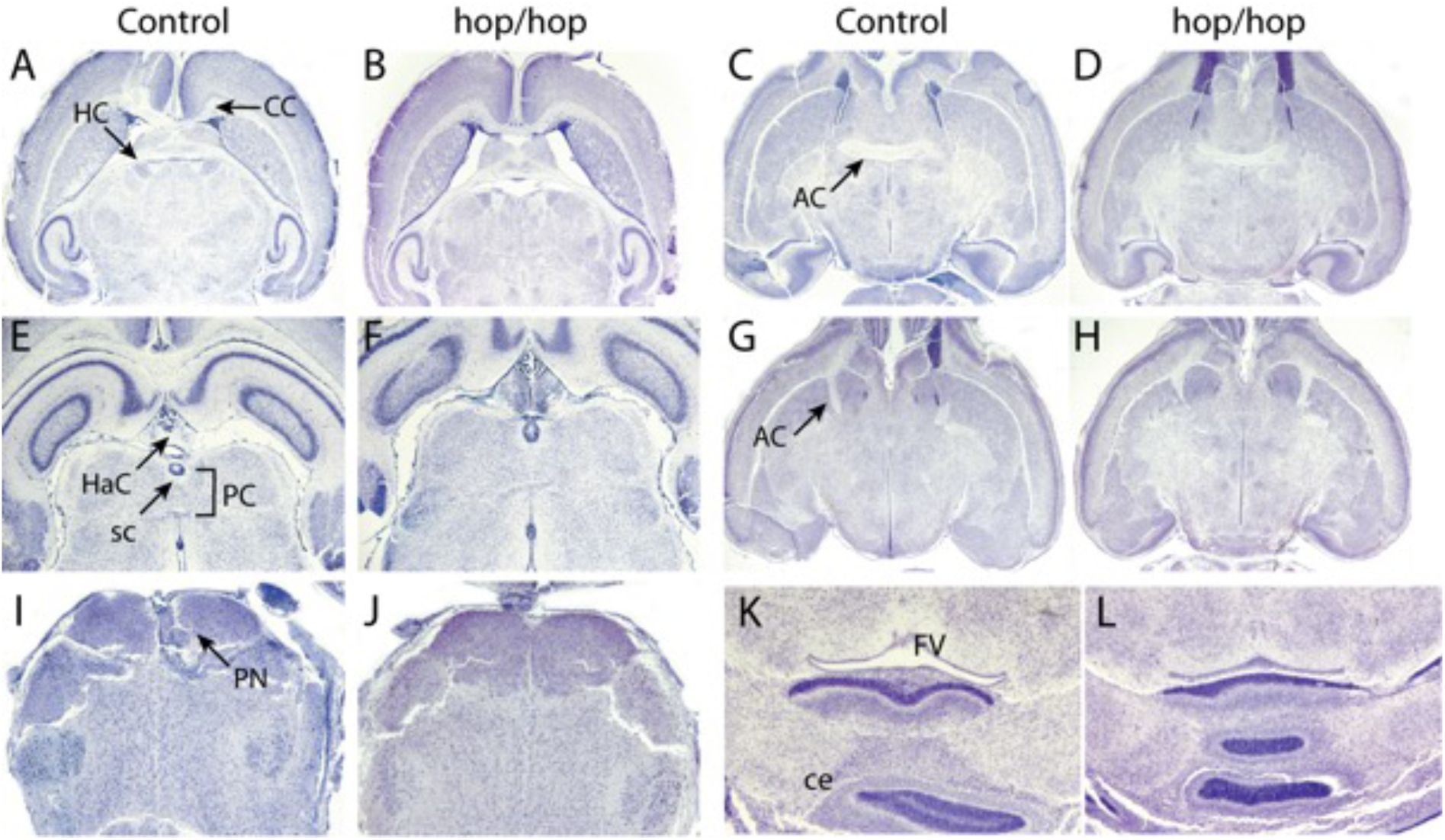
Brain commissure morphology appears normal in *hop* mice. Photomicrographs of horizontal sections from adult control (A,C,E,G,I,L) and *hop* (B,D,F,H,J,L) tissue. (A,B) The corpus callosum (CC) and hippocampal commissure (HC) showed no gross defects. (C,D,G,H) The anterior commissure (AC) also appeared normal. (E,F) Sections showing the habenular commissure (HaC), the posterior commissure (PC) and the subcommissural organ (sc), all three present and with normal appearance in both control and *hop* mice. (I,J) The pontine nuclei (pn) is also present in both genotypes. (K,L) No aberrant commissure was found between the roof of the fourth ventricle (FV), and cerebellum (ce) at the junction of midbrain and hindbrain, in control and *hop* mice.

**Figure S4:**
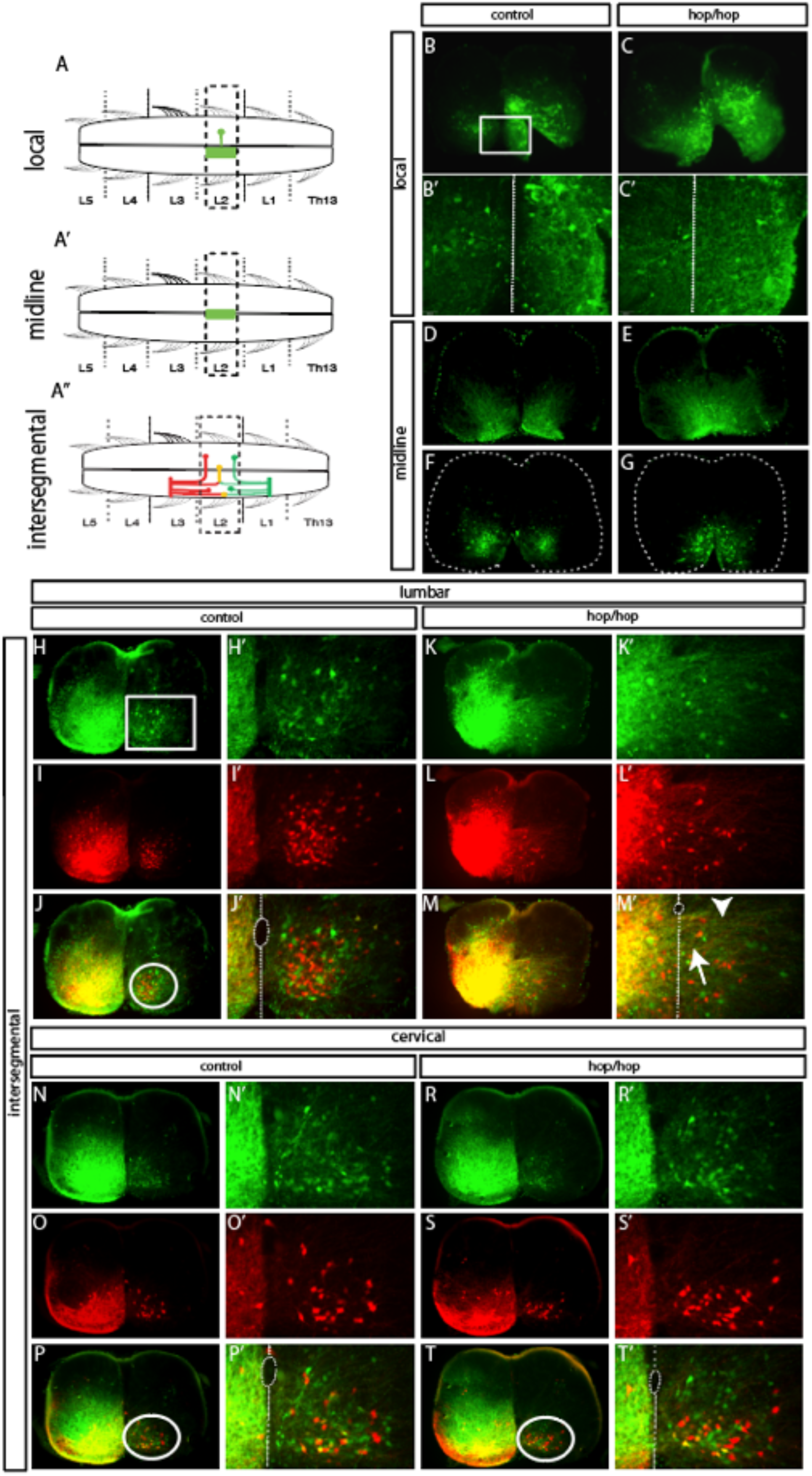
Aberrant axon midline crossing in hop mice. (A-A’’) Schematic drawing of tracer applications in the ventral spinal cord. (A) midline, (A’) local and (A’’) intesegmental tracing using FDA and FDA/RDA to locate local, ascending (aCINs), descending (dCINs) and bifurcating commissural interneurons (adCINs). Photomicrographs of locally projecting CIN according to A (B-C’) or midline traced CINs (D-G). Photomicrographs of traced lumbar spinal cords (H-M’) and cervical spinal cords (N-T’) according to A’’ with FDA and RDA labelled CIN populations in a double exposure. Abundant aberrant midline crossing fibres in *hop* mice (K-M’) compared to controls (H-J’) on lumbar sections. No apparent defects in *hop* mice (R-T’) compared to controls (N-P’) on cervical sections were found.

**Figure S5:**
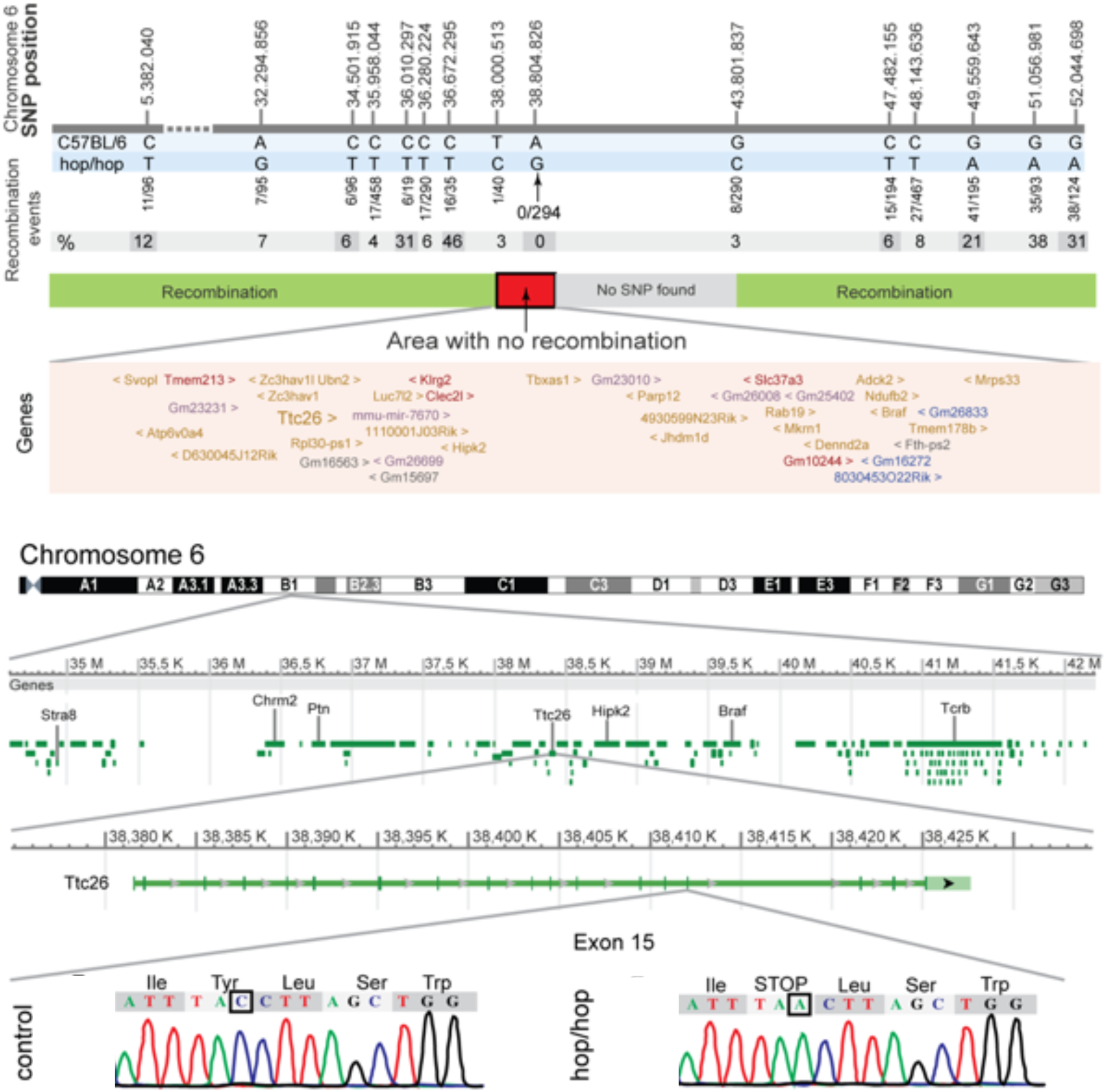
Validation of the *hop* mouse mutation. (A) Region on mouse chromosome 6 harbouring the *hop* locus. SNPs used for linkage mapping are given on the top. The region showing no recombination with *hop* among 800 informative meiosis events is indicated in red. The exon/intron organization of *Ttc26* is indicated together with Sanger sequencing traces of wild type and *hop* highlighting the premature stop mutation in exon 15 (out of 18) of *Ttc26*.

